# An antibody-drug conjugate exploiting a bacterial immune evasion mechanism is effective against multidrug resistant *Neisseria gonorrhoeae*

**DOI:** 10.1101/2025.04.16.644158

**Authors:** Hayley Lavender, Timothy. A. Barendt, Daniel Lucy, Christoph M. Tang

## Abstract

*Neisseria gonorrhoeae* is the causative agent of gonorrhoea, a sexually transmitted infection which is rising in incidence, with increasing drug-resistant strains posing a significant public health threat. To address the urgent need for novel therapies, we developed an antibody–drug conjugate (ADC) that targets this important human pathogen. We utilised Tridecaptin A_1_ a potent antimicrobial peptide against Gram-negative bacteria but exhibits significant toxicity against human cells, limiting its development for clinical use. By conjugating the Tridecaptin A_1_ analogue, Oct-TriA_1_ to a monoclonal antibody (mAb) that specifically targets gonococcal MtrE, the outer membrane component of a drug efflux pump that is upregulated in resistant strains, we aim to deliver selectively deliver the antimicrobial peptide to the gonococcus. However, Oct-TriA_1_ was not bactericidal when directly conjugated to mAb. To circumvent this, we exploited an immune evasion mechanism employed by the gonococcus by introducing a linker between Oct-TriA_1_ and the mAb which is specifically cleaved by the IgA protease (IgAP) secreted by the gonococcus; the IgAP inactivates human IgA. This ADC has no detectable toxicity for relevant human cells, kills the gonococcus in an MtrE- and IgAP-dependent manner, and is active against a strain which is resistant to first line agents. This modular ADC platform could be extended to other bacterial pathogens which employ proteases that use proteases to evade immune killing, offering a new strategy in the fight against antimicrobial resistance.

## INTRODUCTION

*Neisseria gonorrhoeae* is an obligate human pathogen, which causes the sexually transmitted infection (STI), gonorrhoea^1^. There are estimated to be over 82 million cases of gonococcal infection worldwide every year with the highest incidence found in low- and middle-income countries^2^. Complications of gonococcal infection disproportionally affect women, and include pelvic inflammatory disease, ectopic pregnancy, and infertility, as well as increased transmission and acquisition of HIV^3-5^. Due to its exquisite adaptation to survival in its host, *N. gonorrhoeae* has evolved sophisticated mechanisms to subvert human immune defence^6-8^, and secretes an IgA protease (IgAP) which specifically cleaves the hinge region of human IgA1 (hIgA1)^9^.

Due to the rapid emergence of antimicrobial resistance (AMR), the World Health Organization has declared *N. gonorrhoeae* as a priority pathogen for the development of novel interventions^10^, while the Centers for Disease Control declared drug-resistant gonorrhoea as an urgent threat^11^. Of particular concern are multi-drug resistant (MDR) strains which are resistant against the first line agents, extended spectrum cephalosporins (ESCs) and azithromycin. Expression of multi-drug efflux pumps is an important mechanism conferring AMR in *N. gonorrhoeae*. The MtrCDE tripartite pump belongs to the resistance-nodulation-cell division family^12,13^, and can export multiple antibiotics from the gonococcus. Trimeric MtrE forms an outer membrane export channel with a periplasmic α-barrel^14,15^, which is linked by periplasmic MtrC to MtrD at the inner membrane. MtrE also acts as the outer membrane channel for other export pumps including the FarAB and MacAB systems^16,17^. Resistance against several antibiotics is enhanced by increased expression of MtrE, either by loss-of-function mutations of *mtrR* which encodes the repressor of the *mtrCDE* operon or *mtrCDE* promotor mutations^18-23^. Therefore, MtrE is an attractive surface-expressed target for therapeutic interventions as it is overexpressed by many drug-resistant isolates^14,24^.

The rapid acquisition of AMR in Gram-negative bacteria such as the gonococcus and the relative lack of novel antibiotics in development has led to renewed interest in the use of antimicrobial peptides (AMPs)^25,26^. Tridecaptins are acylated tridecapeptides produced by *Paenibacillus* and *Bacillus* spp..^27^ Tridecaptin A_1_ (TriA_1_) is highly active against Gram-negative bacteria although its activity against *N. gonorrhoeae* is unknown^28^. Tridecaptins disrupt cell membranes with its acyl chain critical for activity. However, due to its amphiphilic nature, TriA_1_ interacts non-specifically with cell membranes through electrostatic interactions^29-31^, leading to significant host cell cytotoxicity which has limited the clinical development of tridecaptins and other AMPs with membrane-disrupting proterties^31^.

Here we describe the development of novel anti-*N. gonorrhoeae* <u>a</u>ntibody–<u>d</u>rug <u>c</u>onjugate (ADC) consisting of a monoclonal antibody (mAb) targeting MtrE which is covalently coupled to the highly active TriA_1_ analogue, Oct-TriA_1_. ADCs have been developed against *Staphylococcus aureus* and *Pseudomonas aeruginosa* in which release of the drug from the antibody is mediated by intracellular host proteases, such as cathepsins^32,33^. These enzymes are only active in particular cellular compartments, limiting the distribution of activity of these ADCs. Instead, we exploited the specific activity of a gonococcal enzyme, the IgAP, to release Oct-TriA_1_ from the mAb targeting MtrE. Thus, the delivery of Oct-TriA_1_ to its intended target, the gonococcal outer membrane, is promoted initially by the ADC binding to MtrE, then by the local release of the AMP mediated by bacterial IgAP. We demonstrate that our ADC, which has an IgAP-cleavable linker between the mAb and tridecaptin, displays MtrE- and IgAP-dependent antimicrobial activity against the gonococcus, and kills an MDR *N. gonorrhoeae* isolate while having no demonstrable host cell toxicity. These modular ADCs offer a flexible and customisable approach to deliver a range of antimicrobial compounds to other AMR bacterial pathogens that secrete proteases that cleave defined linear sequences.

## RESULTS

### Oct-TriA_1_ has potent antimicrobial activity against N. gonorrhoeae

The tridecaptin analogue, Oct-TriA_1_ has potent bactericidal activity against a range of Gram-negative bacteria including *Klebsiella pneumoniae* and *Acinetobacter baumannii^30^*. To establish whether Oct-Tri_1_ is also active against *N. gonorrhoeae*, we determined its minimum inhibitory concentration (MIC) against eight *N. gonorrhoeae* WHO reference strains with known antibiotic susceptibility profiles^34,35^, as well as the well-characterised isolates, FA1090 and MS11^34,36,37^. Oct-Tri_1_ displays marked anti-gonococcal activity with MICs in the sub-μg ml^-1^ range for all isolates (0.2-0.78 μg ml^-1^); Oct-Tri_1_ is also active (MIC, 0.78 μg ml^-1^) against isolate G97687, a MDR isolate which is resistant to ESCs and azithromycin^38^ [**Table 1**]. Additionally, the MICs for FA1090Δ*mtrE* (0.1 and 0.2 μg ml^-1^ for the *mtrE* mutant and wild-type, respectively) and MS11Δ*mtrE* (0.2 and 0.78 μg ml^-1^, *mtrE* mutant and wild-type, respectively) were lower than the corresponding wild-type strains, suggesting that Oct-TriA_1_ is a substrate of MtrE-dependent efflux pumps.

**Table 1.** Comparison of the antimicrobial activity of Oct-TriA1 with different *N. gonorrhoea* isolates ^†^.

| Peptide | Neisseria gonorrhoea Strain |  |  |  |  |  |  |  |  |  |  |  |  |  |  |
| --- | --- | --- | --- | --- | --- | --- | --- | --- | --- | --- | --- | --- | --- | --- | --- |
|  | WHO F | WHO G | WHO K | WHO L | WHO M | WHO N | WHO O | WHO P | MS11 | MS11ΔmtrE | MS11ΔIgAP | FA1090 | FA1090ΔmtrE | FA1090ΔIgAP | G97687 |
| Oct-TriA<sub>1</sub> | 0.2 | 0.39 | 0.78 | 0.2 | 0.39 | 0.39 | 0.39 | 0.39 | 0.78 | 0.2 | 0.39 | 0.2 | 0.1 | 0.1 | 0.2 |
| Oct-TriA<sub>1</sub>-c | 1.56 | 3.13 | 3.13 | 1.56 | 1.56 | 3.13 | 3.13 | 3.13 | 3.13 | 1.56 | 6.25 | 0.78 | 0.78 | 0.78 | 1.56 |

†Activities are reported in µg/ ml

### An ADC with Oct-TriA_1_ directly conjugated to an α-MtrE mAb lacks activity against N. gonorrhoeae

The use of tridecaptins for treating Gram-negative pathogens has been hampered by significant host cell toxicity^31^. Therefore, we decided to conjugate an Oct-TriA_1_ analogue to a mAb that binds a gonococcal surface protein, thereby targeting the AMP to *N. gonorrhoeae*. We generated mAbs against MtrE, the surface component of the several gonococcal export pumps^39,40^, because MtrE is overexpressed by many drug-resistant *N. gonorrhoeae* isolates including G97687^38,41^. We examined the conservation of MtrE in *N. gonorrhoeae* and identified 94 unique protein sequences in over 18,600 strains in PubMLST BIGSdb^42^. The five most prevalent MtrE sequences account for >96% of gonococcal isolates [**Figure 1a**, **Extended data Table 1**]. The location of polymorphic residues in MtrE was mapped onto its structure (PDB:4MT0)^15^ [**Figure 1b**], and revealed that the two surface exposed loops of MtrE are highly conserved; a single sequence of loop 1 (GSLSGGNV) and loop 2 (SVELGGLFKSGTG) are found in 99.9% and 98.2% of strains, respectively [**Extended data Table 2**]; most polymorphisms are located in the periplasmic α-barrel of MtrE. This makes MtrE an excellent candidate for targeting the gonococcus.

**Figure 1.**
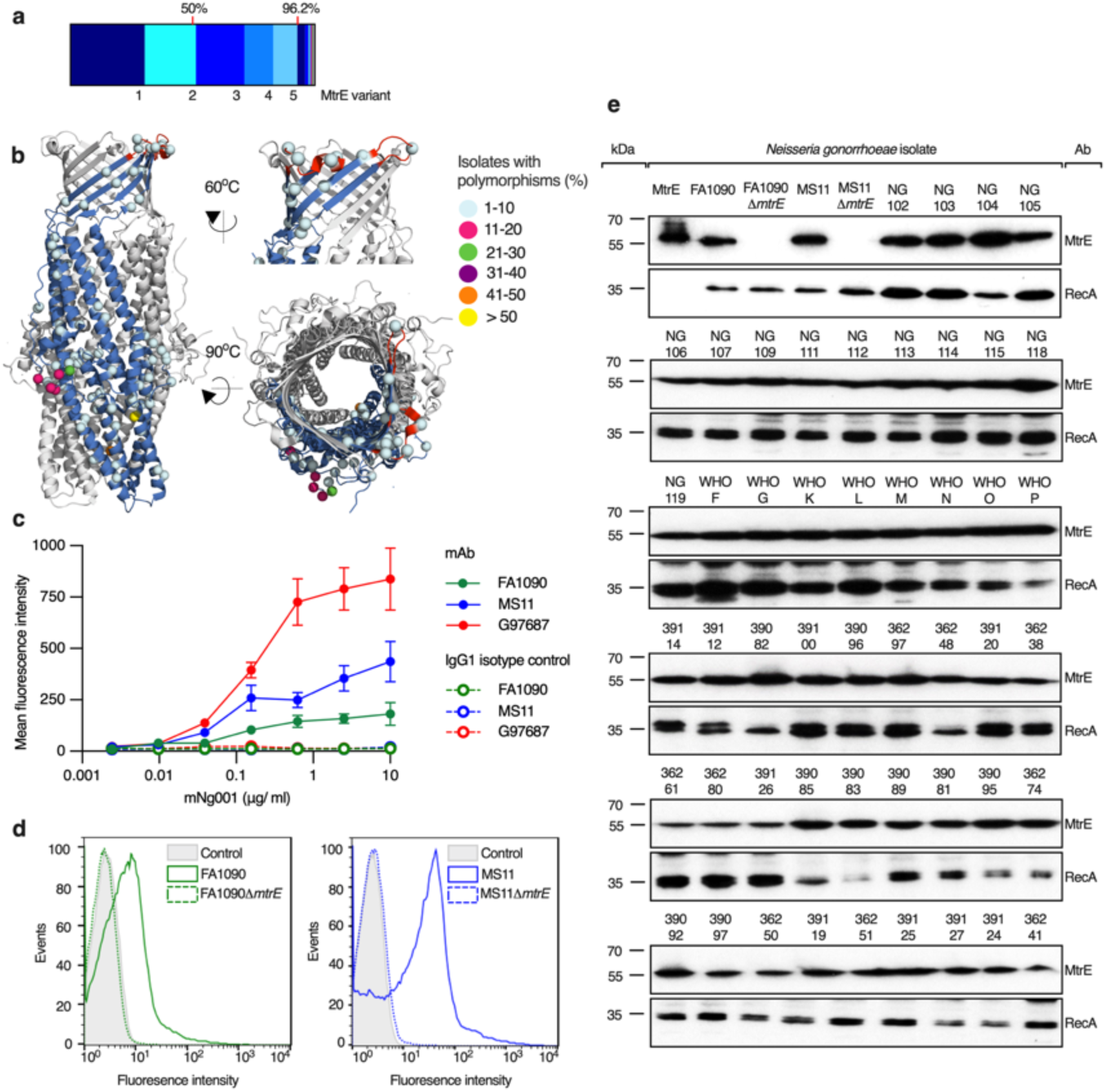
mAb mNg001 recognises gonococcal MtrE. **(a)** Distribution of MtrE amino acid sequences in *N. gonorrhoeae* isolates on PubMLST (n = 18,635). Percentage of isolates harbouring MtrE variants represented as Parts of a Whole. Red lines indicate 50% and 96.2% of *N. gonorrhoeae* isolates. **(b)** Amino acids coloured according to the proportion of isolates with polymorphisms compared to the most common MtrE variant (*i.e.* encoded by allele 1), and mapped onto trimeric MtrE (PDB accession number 4MT0); Outer membrane loops (red) **(c)** Dose-dependent binding of mAb mNg001 (filled circles), or isotype control antibody (unfilled circles), to *N. gonorrhoeae* strains FA1090 (green circles), MS11 (blue circles) and G97687 (red circles). Mean florescence intensity ± standard deviation determined by flow cytometry (n = 3). **(d)** mNg001 binds to wild-type strains; FA1090 and MS11 (solid lines) but not the isogenic Δ*mtrE* mutants (dashed lines). **(e)** Western blot analysis of mNg001 binding to MtrE in whole cell lysates of 52 *N. gonorrhoeae* strains selected from a variety of core genome groups. Recombinant MtrE and wild-type FA1090 and MS11 and Δ*mtrE* mutants used as mAb controls; RecA, loading control.

mAbs were generated against recombinant MtrE by standard hybridoma technology. We characterized one mAb, mNg001, in more detail. Flow cytometry demonstrated that mNg001 binds *N. gonorrhoeae* in a dose-dependent manner [**Figure 1c**] and exhibited higher binding to G97687 and MS11 compared to FA1090, consistent with MtrE overexpression by the former two strains^13,38^. As expected, mNg001 binding was MtrE-specific as no binding was detected to FA1090Δ*mtrE* or MS11Δ*mtrE* (p= 0.0003 and 0.0018, respectively, compared to wild-type bacteria [**Figure 1d**]). We also assessed the ability of mNg001 to recognise MtrE from 52 gonococcal strains from different core genome groups^43^ and included the five most prevalent MtrE alleles by western blot analysis [**Extended data Table 1**]; mNg001 bound to all gonococcal strains analysed [**Figure 1e**]. In contrast, the mAb did not recognise any protein in seven *Neisseria meningitidis* isolates or 16 isolates of other *Neisseria* spp. [**Extended data Figure 1**]. Sequence analysis [**Extended data Figure 2**] shows that meningococcal and commensal Neisseria MtrE share 95.7%-97.5% amino acid identity to the most common gonococcal MtrE variant. Six amino acids are consistently different in MtrE from *N. meningitidis* and commensals compared with the gonococcus [**Extended data Figure 3**]. Five substitutions (Q39, S43, S46, R248, A334S) are located in the α-helical periplasmic domain and β-barrel of MtrE whereas there is only one substitution, V320A, within the outer membrane loop 2 which may explain the species specificity of mAb mNg001. Taken together, mNg001 binds a conserved epitope present in MtrE from *N. gonorrhoeae* but not other *Neisseria* spp., making it suitable for specific delivery of AMPs.

Next, we examined sites in Oct-TriA_1_ that would allow conjugation to mNg001. Oct-TriA_1_ depends on its N-acyl chain and several amino acids, while the C-terminal alanine is dispensable^29^. Therefore, we reasoned that this tridecaptin might retain its activity when directly conjugated directly to mAb mNg001 *via* its C terminus. We covalently conjugated mNg001 to Oct-TriA_1_ with two 8-amino-3,6-dioxaoctanoic acid [O2Oc] spacer groups and a cysteine at the C terminus; *m*-maleimidobenzoyl-*N*-hydoxysuccinimide (MBS) was used as the crosslinker to generate amine-sulfhydryl linkages^44^ [**Figure 2a**]. This ADC was analysed by matrix assisted laser desorption/ionization time-of-flight mass spectrometry (MALDI-TOF MS), which demonstrated that the average drug-to-antibody ratio (DAR) was approximately 10 [**Figure 2b**]. However, the ADC lacked any detectable bactericidal activity against *N. gonorrhoeae* even at concentrations of >512 µg ml^-1^ (equivalent of 70 µg ml^-1^ of Oct-TriA_1_), indicating that Oct-TriA_1_ is inactive when bound to a mAb.

**Figure 2.**
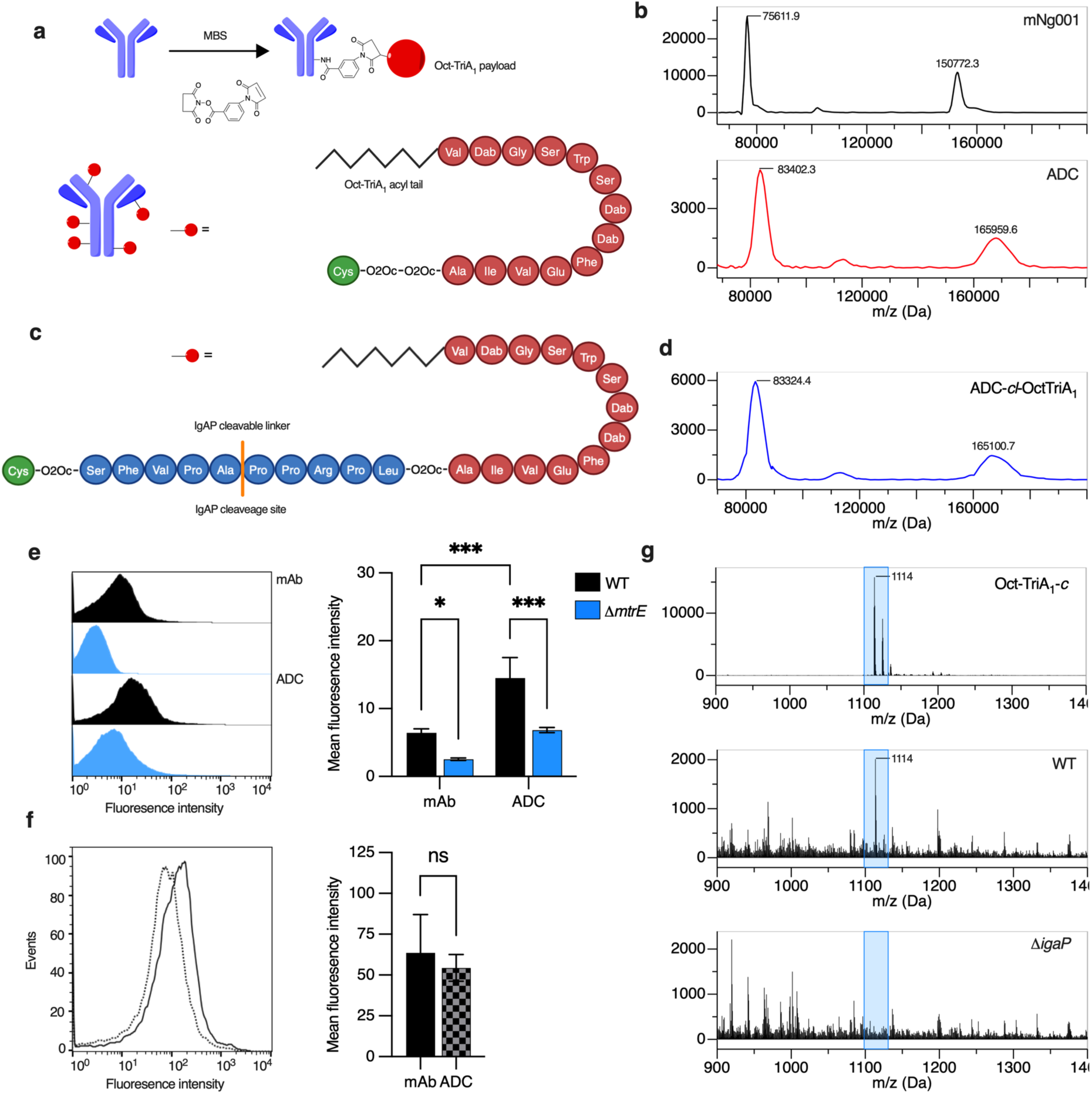
Construction and characterization of ADC with an IgAP cleavable linker. **(a)** (Top) Conjugation of mNg001 (purple) to OctTriA_1_ (Red circle) with MBS containing N-hydroxysuccinimide (NHS) ester and maleimide groups. (Bottom) Model of directly conjugated ADC (not drawn to scale) consisting of mNg001 (purple) and OctTriA_1_ analogue (red) synthesized with two PEG-based linkers (O2Oc, 8-amino-3,6-dioxaoctanoic acid) and a C-terminal cysteine (green), acyl tail (black). Dab = diaminobutyric acid **(b)** MALDI-TOF analysis of mNg001 (top, black) and directly conjugated ADC covalently linked to synthesized OctTriA_1_ analogue (red). Change in mass, compared to mNg001 indicates a DAR of approximately 10 peptides per ADC. **(c)** Schematic of *cl*-OctTriA_1_ analogue incorporating an IgAP cleavable linker [LPRPPAPVFS] between two 8-amino-3,6-dioxaoctanoic acid linkers with a C-terminal cysteine. **(d)** MALDI-TOF mass spectrometry of *cl*-ADC indicates a DAR of approximately 5-6 peptides. Binding of mNg001 (mAb) and ADC conjugated to *cl*-OctTriA_1_ to gonococcal strains **(e)** FA1090 wild-type (black) and isogenic Δ*mtrE* (blue) or **(f)** G97687. Data show mean ± standard deviation (n = 3). two-way ANOVA; *** p < 0.001, * p ≤ 0.05 and ns, p > 0.05. **(g)** Mass spectrometry analysis of OctTriA_1_-*c* (blue box) alone (top), and the ADC after incubation in supernatants from wild-type FA1090 (middle) but not FA1090Δ*igaP* (bottom). Synthesised OctTriA_1_-*c* in PBS shown as a mass spectrometry control.

### Introduction of an IgAP cleavable linker into an ADC allows tridecaptin release

Therefore, we decided to exploit the activity of the IgAP secreted by gonococci to release Oct-TriA_1_ from the α-MtrE mAb. Of note, non-invasive *Neisseria* spp., in contrast to *N. gonorrhoeae* and *N. meningitidis*, do not secrete an IgAP^9^ adding another layer of specificity to the ADC [**Extended data Table 3**]. The gonococcal IgAP recognises and cleaves linear 10 amino acid peptides^45^. Therefore, we synthesised Oct-TriA_1_ with an IgAP-cleavable linker (LPRPP/APVFS, / indicates the cleavage site) flanked by O2Oc groups to the C terminus of Oct-TriA_1_; a C-terminal cysteine was added after the linker (generating a <u>cl</u>eavable Oct-TriA_1_, cl-Oct-TriA_1_) to allow conjugation to the mAb [**Supplementary table 1, Figure 2c**].

We examined whether cl-Oct-TriA_1_ can be cleaved by the gonococcal IgAP. cl-Oct-TriA_1_ was incubated with culture supernatants from wild-type FA1090 or an isogenic Δ*igaP* mutant. Electrospray ionisation mass spectrometry (ESI-MS) analysis of the uncleaved peptide showed species with a mass-to-charge ratio (m/z) corresponding to *cl*-Oct-TriA_1_, of 1488 [**Extended data figure 4**]. After incubation with supernatants from wild-type bacteria but not from the Δ*igAP* mutant, a species with a m/z of 1114 [**Extended data figure 4**] was detected from cl-Oct-TriA_1_, consistent with the mass of cleaved Oct-TriA_1_ (*i.e.,* Oct-TriA_1_ with -O2Oc-LPRPP). To determine whether cleaved cl-Oct-TriA_1_retained its activity, we synthesised a peptide corresponding to cl-Oct-TriA_1_ after cleavage by the IgAP (Oct-TriA_1_-*c*) [**Supplementary table 1**] and found that it is highly active against *N. gonorrhoeae* (MICs, 0.78 to 6.25 μg ml^-1^, [**Table 1**]).

As gonococcal IgAP can cleave cl-Oct-TriA_1_ and the residual AMP retains bactericidal activity, we next determined if the gonococcal IgAP could cleave cl-Oct-TriA_1_ conjugated to mAb mNg001. Again, we used MBS to covalently link cl-Oct-TriA_1_ to the mAb [**Figure 2c**]. The ADC was subjected to MALDI-TOF MS which demonstrated a DAR of ∼6 [**Figure 2d**]. We also generated ADCs conjugated to an identical Oct-TriA_1_ analogue but with an <u>uncl</u>eavable 10 amino acid linker (uncl-Oct-TriA_1_, with VKPAPSPAAN) [**Supplementary table 1**] uncl-Oct-TriA_1_, which also yielded a DAR of ∼6 [**Extended data figure 5**]. To assess if the ADCs still recognise *N. gonorrhoeae* after conjugation, we analysed the binding to FA1090 of mAb mNg001 or the ADCs. Flow cytometry demonstrated that binding of the ADCs was dependent on MtrE expression [**Figure 2e**] with significantly decreased binding to FA1090Δ*mtrE* compared to FA1090 (p= 0.0003). The ADC displayed marginally more binding to FA1090 and FA1090Δ*mtrE* than the unconjugated mAb (p= 0.0002 and 0.0093 respectively, [**Figure 2e**] which may indicate low level, non-specific electrostatic interactions with the AMP^29-31^. Furthermore, we compared binding of mNg001 and the ADCs to the multidrug resistant isolate G97687, and no significant difference in binding was observed to this strain (p= 0.5586, [**Figure 2f**]).

To examine if the IgAP can release Oct-TriA_1_ from the mAb, we incubated the ADCs with supernatants from wild-type FA1090 or the *ΔigAP* mutant. ESI-MS confirmed that the presence of IgAP liberates Oct-TriA_1_ from the ADC generated with cl-Oct-TriA_1_ (a species with m/z of 1114 m/z corresponding to free, cleaved cl-Oct-TriA_1_, Oct-TriA_1_-*c*, **Figure 2g**) but not the ADC generated with uncl-Oct-TriA_1_ (*i.e.,* with the non-cleavable linker), demonstrating that the gonococcal IgAP specifically liberates Oct-TriA_1_ from ADCs containing the cleavable linker [**Extended data Figure 6**].

### ADCs exhibit reduced host cytotoxicity

Tridecaptins cause significant host cell cytotoxicity^29^. Therefore, we next evaluated the hemolytic activity (HC_50_) of Oct-TriA_1_ derivatives against human erythrocytes [**Figure 3a**]. Unmodified Oct-TriA_1_ had a HC_50_ of 39.4 μg ml^-1^, while Oct-TriA_1_-*c* (the peptide generated by IgAP cleavage of the ADC) demonstrated >25-fold reduced hemolytic activity (HC_50_ of >1 mg ml^-1^). Of note,the HC_50_ of the ADCs was > 2.8 mg ml^−1^, equivalent to > 400 μg ml^−1^ of Oct-TriA_1_-*c* with a DAR of 8 [**Figure 3b**], demonstrating that conjugation of Oct-TriA_1_ to a mAb virtually abolishes toxicity.

**Figure 3.**
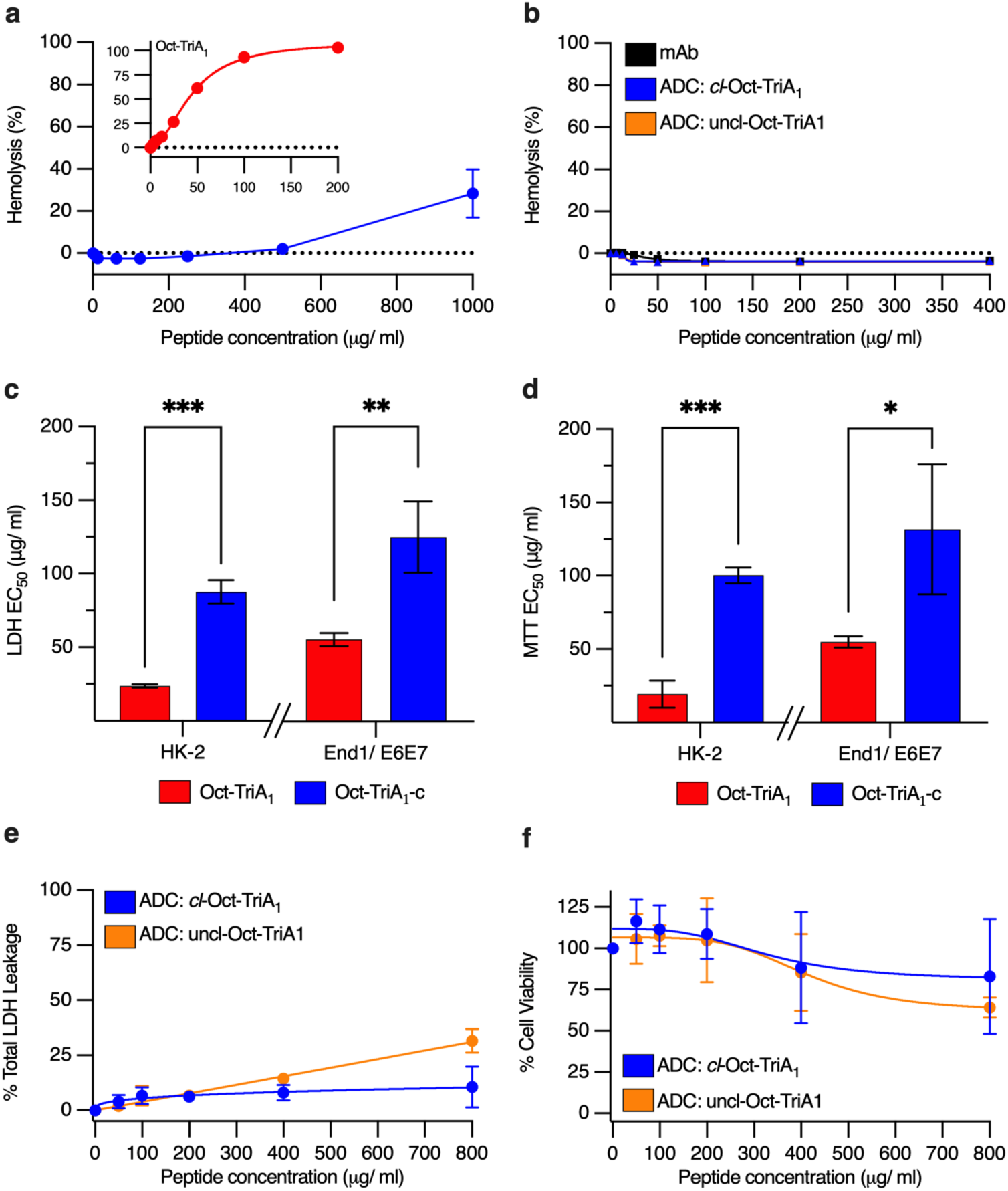
Conjugation of OctTriA_1_ to ADC reduces host cell cytotoxicity. Hemolysis of human erythrocytes by OctTriA_1_ derivatives. **(a)** OctTriA1, (red, inset), OctTriA_1_-*c* (blue) and **(b)** mNg001 (black) with ADCs conjugated to *cl*-OctTriA_1_ (blue) and un*cl*-OctTriA_1_ (orange). Data are the mean ± standard deviation (n = 3). Cytotoxicity of OctTriA_1_ peptides against HK-2 renal proximal tubular and End1 E6/E7 endocervical cells measured by **(c)** LDH release and **(d)** reduction of MTT; OctTriA1, (red) and OctTriA_1_-*c* (blue). Cytotoxicity of ADCs conjugated to *cl*-OctTriA_1_ (blue) and un*cl*-OctTriA_1_ (orange) towards HK-2 cells assayed by **(e)** LDH release and **(f**) MTT reduction. Data are the mean ± standard deviation (n = 3). Unpaired *t* test; *** p < 0.001, ** p <0.01, and *p ≤ 0.05.

Additionally, we assessed the toxicity of Oct-TriA_1_ derivatives against HK-2 human proximal renal tubule cells^46^, and upper cervical End1/E6E7 cells^47^ using LDH and MTT assays. These cell lines were chosen as AMPs usually cause nephrotoxicity while the gonococcus infects cells in the upper vagina and cervix. The half-maximal inhibitory concentration (IC_50_) for Oct-TriA_1_-*c* was significantly higher than Oct-TriA_1_ for both the LDH release (p= 0.0002 and 0.0081) and cell viability (p= 0.0002 and 0.0406) with HK-2 and End1/E6E7 cells, respectively [**Figure 3c-d**, **Table 2**]; End1/E6E7 cells have increased tolerance for Oct-TriA_1_ analogues, demonstrated by the higher IC_50_ for Oct-TriA_1_-*c* than HK-2 cells [**Table 2**]. Remarkably, the ADCs had an IC_50_ > 5.6 mg ml^−1^, equivalent to > 800 μg ml^−1^of Oct-TriA_1_ [**Figure 3e-f**, **Table 2**]. Taken together, modification of the C-terminus of Oct-TriA_1_ significantly reduces host-cell toxicity, while the ADCs displayed no detectable toxicity against cells from the renal and genitourinary tracts.

**Table 2.**
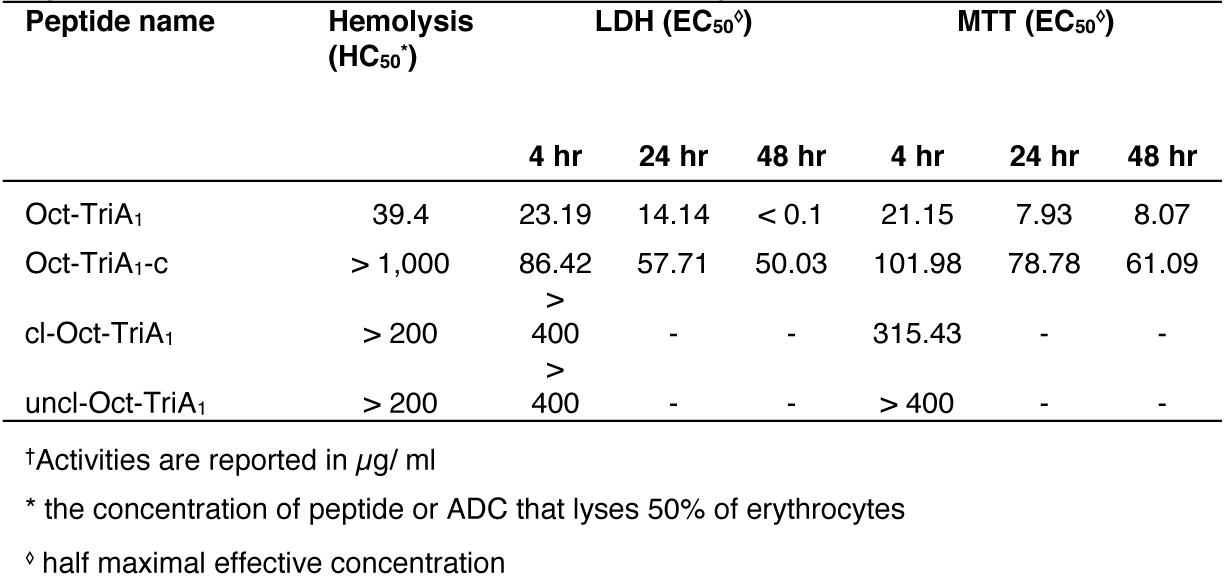
Comparison of human erythrocyte hemolysis (HC50) and cytoxicity (EC50) against HK-2 cells on exposure to Oct-TriA1 analogues and ADCss †.

### ADCs have MtrE- and IgAP-dependent anti-gonococcal activity and are active against an MDR strain

Finally, we assessed the bactericidal activity of the ADCs against *N. gonorrhoeae*. As bacteria need to secrete IgAP for to release the antimicrobial peptide from ADCs, we assessed the activity of ADCs in time-kill assays rather than MICs [**Figure 4**]. Oct-TriA_1_-*c* exhibited dose-dependent bactericidal activity against FA1090 and the MDR strain G97687 in time-kill assays, with undetectable survival of FA1090 in 50 μg ml^-1^ of the AMP after 60 min; there was no survival of G97687 after 180 minutes [**Figure 4a-b**]; G97687 has a higher MIC than FA1090 for Oct-TriA_1_-*c* [**Table 1**].

**Figure 4:**
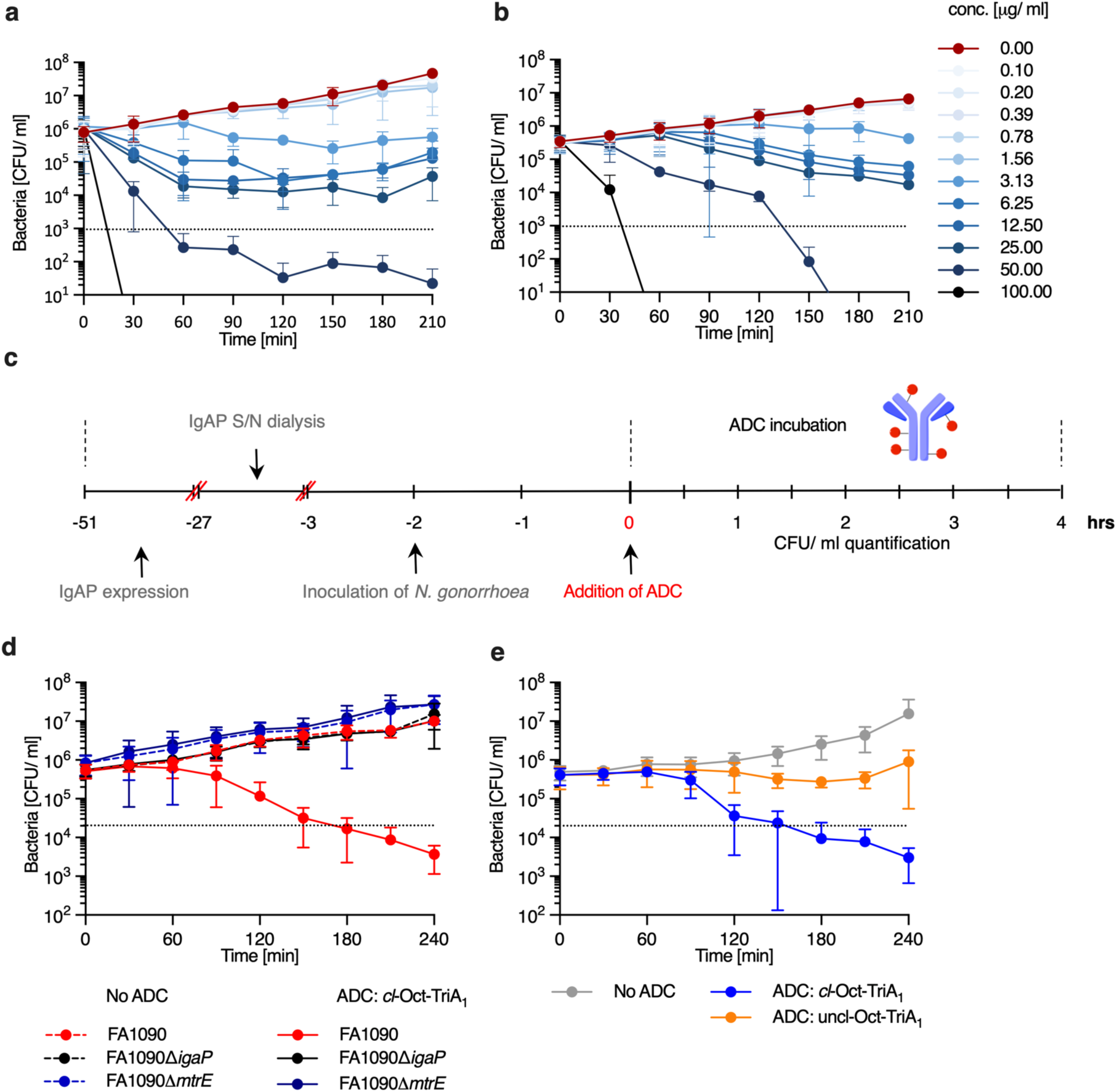
ADCs are bactericidal in the presence of the gonococcal IgAP. Dose-dependent time to kill analysis of OctTriA_1_-c against **(a)** FA1090 and **(b)** G97687. No peptide is shown in red with increasing concentrations of peptide from 0.1 μg/ ml (light blue) to 100 μg/ ml (dark blue). **(c)** Time to kill curve analysis with ADCs protocol. *N. gonorrhoeae* were grown for 24 hours prior to harvesting supernatant with or without IgAP, wild-type and Δ*igAP* respectively. Supernatant was dialysed into fresh media prior to inoculating with respective *N. gonorrhoeae* strains. After 3 hours growth, ADCs or peptides were added, and CFU/ ml was calculated every 30 minutes for 4 hours. **(d)** Time to kill curve analysis of ADC-*cl*-OctTriA_1_ against FA1090 (red), isogenic *ΔIgAP* (black) and *ΔMtrE* (blue). No ADC (dashed line) and ADC (line). **(e)** Time to kill curve analysis of ADC-*cl*-OctTriA_1_ (blue), ADC-un*cl*-OctTriA_1_ (orange) and mNg001 (grey) against the MDR gonococcal strain, G97687. All data shown are the mean ± standard deviation of CFU/ml (n = 3).

As each ADC has an average DAR of 7-8 we used a concentration equivalent to 100 μg ml^−1^ of Oct-TriA_1_ in time-kill assays; initial bacterial growth allowed expression of the IgAP prior to treatment with ADC’s [**Figure 4c**]. The survival of FA1090 decreased after 90 min incubation with the ADC containing Oct-TriA_1_ conjugated *via* the cleavable linker and was below the limit of detection (LOD) after 180 min [**Figure 4d**]. In contrast, the ADC with Oct-TriA_1_ conjugated *via* a non-cleavable linker had no bactericidal activity against FA1090 [**Extended data figure 7**]. The ADC with the cleavable linker displayed no bactericidal activity In the absence of MtrE, the mAb target (*i.e.* against FA1090Δ*mtrE*) or the IgAP (*i.e*. against FA1090Δ*igaP*). Importantly, this ADC efficiently killed the MDR isolate G97687 [**Figure 4e**]. Taken together, our data demonstrate that Oct-TriA_1_ conjugated to an α-MtrE mAb *via* a IgAP cleavable linker offers targeted delivery of an antimicrobial peptide that efficiently kills MDR *N. gonorrhoeae* while reducing off-target cytotoxicity. The mechanism of action of this ADC is dependent on both its ability to bind MtrE on the bacterial surface, and the IgA protease secreted by the gonococcus as an immune evasion strategy.

## DISCUSSION

*N. gonorrhoeae* is an urgent public health treat due to its increasing incidence and increasing AMR^2^. Resistance has emerged in the gonococcus within a few years of introduction of every antibiotic^48^. The rise in AMR not only undermines the treatment of patients with gonococcal disease but also jeopardises the control of infection, which relies on successful treatment of their contacts^38,49^. Post-exposure prophylaxis (PEP) is being used to protect individuals who have potentially been exposed to STIs auer unprotected intercourse^50^. Efforts to use doxycycline against gonorrhoea have been less successful against the gonococcus than for other STIs due to plasmid-mediated AMR^43,50^. Therefore, there is an urgent need to develop alternative treatment options against this important human pathogen.

Tridecaptins are potent AMPs which we show are highly active against *N. gonorrhoeae*^30^. However, these cationic peptides also interact non-specifically with lipid-containing membranes, leading to unwanted cytotoxicity and hemolytic activity^31^. This inherent toxicity has limited the development of tridecaptins and other AMPs for treating bacterial infection. Instead, we repurposed these potent AMPs by conjugating them to a gonococcal specific mAb to deliver them directly to the bacterial surface. Although ADCs have been extensively used in oncology^51^, only three mAbs have been approved for treating bacterial infection, all of which neutralise secreted toxins^52-54^ rather than having direct antimicrobial activity. Efforts have been made to develop ADCs which are bactericidal against *S. aureus* and *P. aeruginosa* by incorporating a host cathepsin-cleavable linker to release a bactericidal payload^55,56^. Unfortunately, cathepsin is only activated in the phagolysosome so such a strategy would not be effective against extracellular pathogens. Instead, we engineered ADCs with bacterially-cleavable linker to allow release of a highly active AMP in close proximity to the gonococcus. This additional targeting mechanism led to an ADC with virtually no host cell cytotoxicity without compromising bactericidal activity.

The mAb in our ADC targets MtrE, which is conserved and constitutively expressed by *N. gonorrhoeae*^12^. MtrE is the outer membrane channel of three gonococcal efflux pumps, MtrCDE, FarAB and MacAB, which contribute to AMR^57^. Importantly, the MtrCDE system is overexpressed by many MDR gonococcal strains^13,38^. The two surface exposed loops of MtrE are highly conserved and our mAb recognises all 51 isolates tested to date, spanning the genetic diversity of the gonococcus^43,58^. To couple Oct-TriA_1_ to mNGO001 we investigated the linear peptide, LPRPPAPVFS, as a bacterial-cleavable linker containing an IgAP cleavage site^45^. The IgAP is a sophisticated immune evasion mechanism deployed by the gonococcus which specifically cleaves hIgA1^9^. Therefore, mutants which escape killing by the ADC by not expressing MtrE or the IgAP will be more susceptible to other antibiotics or local immune responses, respectively.

To synthesize the ADC, we identified a site on Oct-TriA_1_ for its ayachment to the mAb. The length of the acyl tail and several residues (*i.e.* D-Trp5, D-Dab^8^ and D-*allo*-Ile^12^) are critical for the antimicrobial activity of Oct-TriA_1_^31^. Glu^10^ of an unacylated tridecaptin analogue has been modified for addition of erythromycin and vancomycin analogues against *Klebsiella pneumoniae*^59^, but the toxicity of these conjugates against host cells is unknown. We found that a 10 amino acid linker (LPRPPAPVFS) can be added to the C-terminal Ala of Oct-TriA_1_ with a negligible effect on antimicrobial activity of the AMP (**Table 1**). cl-Oct-TriA_1_ was covalently conjugated to our gonococcal-specific, α-MtrE mAb by maleimide cross-linking between a C-terminal Cys group on cl-Oct-TriA_1_ and lysine residues on the mAb^60^. This resulted in an average DAR of 7-8.

Time-kill curves demonstrate that the ADCs with a cleavable linker are bactericidal against *N. gonorrhoeae* in an MtrE- and IgAP-dependent manner. Furthermore, the cleavable ADC kills of the highly resistant isolate, G97687, which is resistant to all first line agents^38^. In contrast, ADCs with an uncleavable linker had no bactericidal at lower concentrations (50 μg ml^-1^), but at higher concentrations (150 μg ml^-1^) exhibited non-specific bactericidal activity. NHS-esters form partially stable linkages which can be hydrolysed in aqueous environments with spontaneous release of the AMP^60^. In the future, specific AMP:mAb linkages will be generated by engineering the AMP and mAbs, resulting in a similar yet defined DAR and known sites of conjugation^33,55^.

Our strategy for developing ADCs against *N. gonorrhoeae* is modular, allowing selection of different mAbs, linkers, and AMPs. The mAb we employed is recognises the five most common variants of gonococcal MtrE without binding MtrE from other species of *Neisseria* spp., providing precise targeting of the ADC to the gonococcus. Additionally, commensal *Neisseria* spp. rarely express the IgAP (**Extended data Table 3b**) making it unlikely that Oct-TriA_1_ is inadvertently released from the ADC by this group of bacteria. The meningococcus does express an IgAP related to the gonococcal enzyme so could cleave the ADC although *N. meningitidis* MtrE is not recognised by mNg001; the IgAP from other genera recognise other amino acid sequences^61^. Thus, the mechanism of release of the AMP provides another layer of specific activity for the ADC, which should be active against the gonococcus irrespective of their site in the host as it relies on a constitutively expressed bacterial immune escape mechanism and not a host protease.

In conclusion, our findings indicate that our ADC is a potent antimicrobial against MDR *N. gonorrhoeae* with virtually undetectable host cell cytotoxicity or hemolytic activity. We have exploited two key virulence factors of the gonococcus, MtrE and IgAP, which have important roles in gonococcal AMR and pathogenesis. To our knowledge, this is the first anti-bacterial ADC containing a linker which can be cleaved by a bacterial protease. The ADC should induce minimal disruption of the microbiome, which is an advantage over other small molecule therapeutics, antibiotics, and AMPs. Finally, our ADCs can also be utilised as a plazorm for the development of other ADCs targeting other bacterial species as the ADC plazorm is entirely modular so the components can be exchanged to target other organisms.

## MATERIALS AND METHODS

### mtrE analyses

*mtrE* sequences were downloaded from PubMLST^42^ (https://pubmlst.org/organisms/neisseria-spp, accessed June/July 2023), translated, then aligned by Muscle in Geneious Prime 2023. The number of isolates associated with each single amino acid polymorphism (SAAP) was imported into GraphPad Prism v10.2.2, then mapped onto the structure of MtrE (PDB accession no. 4MT0)^15^.

### Bacterial strains and growth

Gonococcal base (GCB)^62^, Fastidious broth (FB)^63,64^, and Graver and Wade medium (GW)^65^ were prepared as previously, and supplemented with 1% Vitox; 1.5% (w/v) agar (Oxoid) was added for solid media. *N. gonorrhoeae* strains (**Supplementary table 2**) were grown on GCB agar for 16–18 hr at 37°C with 5% CO_2_. For liquid growth, bacteria were inoculated into 125 ml conical flasks containing 20 ml of liquid medium at an OD_A600_ of 0.1, then incubated in 5% CO_2_ at 37°C shaking at 150 r.p.m.; kanamycin was added to media at 80 μg ml^−1^ as required. *Escherichia coli* was grown in Luria-Bertani (LB) broth, Terrific broth (TB), or on LB with 1.5% (w/v) agar. Antibiotics were added at the following concentrations for *E. coli*: kanamycin, 50 μg ml^−1^, and carbenicillin, 100 μg ml^−1^.

### Generation of ΔmtrE and ΔigaP strains

Genomic DNA was isolated using the Wizard Genomic DNA purification kit (Promega), and up- and down-stream fragments of *mtrE* and *igaP* were amplified using Herculase II Fusion DNA Polymerase (Agilent); primers (**Supplementary table 2**) were designed with the NEBuilder® assembly tool. Amplified fragments were inserted into pUC19 flanking a kanamycin resistance cassette using Gibson Assembly® master mix (New England Biolabs) then transformed into *E. coli* DH5α. Plasmids were confirmed by sequencing, digested with *Aat*II and *Bsa*I prior to transforming *N. gonorrhoeae* as previously described^62^.

### Generation of α-MtrE mAbs

*mtrE* was amplified from *N. gonorrhoeae* FA1090 and ligated into pET14b, generating pET14b:*mtrE*. *E. coli* B834 containing pET14b:*mtrE* was grown in TB for 24 h at 22°C and harvested by centrifugation at 5,000 *× g* for 30 min at 4°C prior to lysis using an EmulsiFlex-C5 homogenizer (Avestin, 15,000 lb/in^2^). Lysates were centrifuged at 20,000 *× g* for 30 min at 4°C, and recombinant MtrE was bound to a HisTrap column (GE Healthcare), eluted with 300 mM imidazole, and purified by size exclusion (AKTA, HiLoad 16/600 Superdex® 200 pg column; GE Healthcare). Protein concentrations were estimated using a Nanodrop 2000c {Thermo Scientific).

Female BALB/c mice (6 to 8 weeks old; Charles River, Margate, UK) were immunised with recombinant MtrE (20 μg) adsorbed to aluminium hydroxide (final composition, 0.5 mg ml^−1^Al-OH_3_, 10 mM histidine-HCl) by mixing overnight at 4°C. Antigens were given by the intraperitoneal route on days 0, 21, and 35. Spleens and sera were harvested on day 49. B lymphocytes were fused to NS0 myeloma cells^66^ with polyethylene glycol (PEG 1500, Roche)^67^. Fused cells were plated in RPMI 1640 containing 2 mM L-glutamine, penicillin (100 U ml^−1^), streptomycin (100 μg ml^−1^), and 1% Ultroser G (Pal France). After 24 hr, 2% hypoxanthine aminopterin thymidine (Life technologies) was added to the wells. Hybridomas were screened by ELISA against recombinant MtrE then cloned by limiting dilution^68^. Animal experiments were carried out under protocols approved by the Home Office UK (PPL 30/3194). mAbs were purified from hybridoma supernatants in Gibco® PFHM II serum-free media supplemented with 2% (w/v) cholesterol by protein G affinity chromatography (Pierce, ThermoFisher Scientific). mAb isotypes were determined using a Mouse Monoclonal Antibody Isotyping Test Kit (Bio-Rad).

### SDS-PAGE and Western blot analysis

Bacteria were grown overnight and 10^9^ CFU ml^−1^ suspended in SDS-PAGE loading buffer (100 mM Tris-HCl, pH 6.8, 20 μM β-mercaptoethanol, 4% SDS, 0.2% bromophenol blue, 20% glycerol) prior to separation on 14% polyacrylamide gels. Proteins were transferred to Immobilon P polyvinylidene difluoride membranes (Millipore, USA) using the Trans-Blot SD semi-dry transfer system (Bio-Rad, USA), then blocked in 3% (w/v) skimmed milk-PBS with 0.05% (v/v) Tween 20 overnight and incubated with hybridoma supernatants or anti-MtrE mAb (5 μg ml^−1^). Membranes were washed three times in PBS with 0.05% (v/v) Tween 20, and incubated with goat anti-mouse HRP (P0447, Dako, UK), washed again, and binding detected using ECL (Amersham, USA).

### Flow cytometry

Approximately 10^8^ CFU ml^−1^ of bacteria were re-suspended in mAb (5 μg ml^−1^) for 30 min, washed in wash buffer (0.05% (w/v) BSA/PBS), then incubated for with 2 μg ml^−1^ goat anti-mouse IgG-Alexa fluor^®^ 647 conjugate (A-21235, Life Technologies) at room temperature. Samples were washed three times then fixed in 3% paraformaldehyde for 30 min, washed three times then resuspended in PBS prior to running on a FACSCalibur (BD Biosciences) or BD LSRFortessa™ Cell Analyzer. Results were analysed by calculating the geometric mean fluorescence (FL-4) in FlowJo v10.10 software (Tree Star). Statistical significance was tested using an unpaired Student *t* test or 2-way ANOVA (GraphPad Prism v.10.2.2).

### Conjugation of Oct-TriA_1_ derivatives with mAbs

*m*-maleimidobenzoyl-*N*-hydoxysuccinimide (MBS, 5 mM) was added to purified mAb (0.1 mM) and incubated at room temperature for 60 min. Excess MBS was removed over ZebraTM Spin desalting columns (ThermoFisher scientific). Activated mAb was added to 0.9 mg of Oct-TriA_1_ analogues and incubated at room temperature for 60 min. ADCs were dialysed using slide-A-Lyzer cassette (20K MWCO, ThermoFisher scientific) into PBS overnight at 4°C and filter sterilised.

### MALDI-TOF MS analysis for determination of intact mass

mAbs or ADCs with a concentration of 1 mg/mL were acidified with 1% trifluoroacetic acid and mixed with Super DHB matrix, 1 μl was spotted onto an MTP Anchorchip 600/384. For every mAb, three sample spots were prepared from the same solution using three protocols advised by the manufacturer. MALDI-TOF measurements were performed on a Microflex LRF mass spectrometer, equipped with a 60Hz nitrogen laser at 337nm.

### Cleavage and detection of Oct-TriA_1_ derivatives and ADC activity

IgA1 protease activity was measured against purified hIgA1 (5 μg, Abcam, UK) incubated with supernatants (20 μl) from *N. gonorrhoeae* grown in GW or FB for 4 or 24 hr, respectively, at 37 °C. Products were detected by Western blot analysis using an α-human IgA mAb-HRP (Cat number A4789, Sigma). Supernatants of *N. gonorrhoeae* grown for 24 hr in FB medium were dialysed into PBS (slide-A-Lyzer, 20K MWCO, ThermoFisher scientific). Peptides (50 μg) or ADCs (750 μg) were incubated for 4 hr at 37°C with supernatants. Samples were desalted then analysed by mass spectrometry with a Waters Micromass LCT Mass spectrometer using electrospray ionization (ESI-MS). *N. gonorrhoeae* was resuspended in PBS to an OD_A600_ 1.0, and ̴5 × 10^4^ CFU added to wells of a microdilution plate containing dilutions of antimicrobials in FB. Plates were incubated for 24 hr at 37°C, in 5% CO_2_ then 10 μl aliquots plated onto GCB agar. The MIC was defined as the lowest concentration of an antimicrobial that prevented bacterial growth.

### Time-kill assays

For time-kill assays^69^, bacterial suspensions (10^8^ CFU) were diluted in GW medium, and 30 μl added to 7.5 ml pre-warmed GW medium. A total of 90 μl of the suspension was added per well (Nunclon™ microtiter plates, ThermoFisher scientific), then incubated for 3 h. Next 110 μl of an antimicrobial or media alone was added to each well and incubated for 4 hr. Samples (10 μl) were plated every 30 min in triplicate onto solid media.

### Cytotoxicity assays

For hemolysis^70^, whole human blood (1 ml; K2EDTA; Cambridge Bioscience) was washed in PBS then 0.5% added to an equal volume of peptides or ADCs in 96-well V-bottomed polypropylene plates and incubated at 37°C for 1 hr. Cells were pelleted at 1,000 *× g* for 10 min and the absorbance at OD_415_ measured. Positive (melittin, 2.5 µM, Merck Life Science) and negative (PBS) controls were normalised as 100 and 0% hemolysis, respectively.

HK-2 cells (CRL-2190™, Americab Type Culutre Collection (ATCC)), a proximal renal tubule cell^46^, and End1/E6E7 (CRL-2615™, ATCC) upper cervical epithelial cells^47^ were cultured in Keratinocyte Serum Free Medium (Invitrogen, Life Technologies Ltd.) with human recombinant EGF (5 ng ml^-1^) and bovine pituitary extract (0.05 mg ml^−1^) at 37°C in 5% CO_2_. Cells were seeded in 96-well plates (Corning, Polystyrene) at a density of 2 × 10^4^ cells/well 24 hr prior; Triton-X-100 (0.1%) and Digitonin (30 μg ml^−1^) were used as controls. Lactate dehydrogenase (LDH) release was measured using the Biovision kit; the OD_490_ was assessed in a SpectraMax M5 plate reader.

Cell viability was assessed by 3-(4,5-dimethylthiazol-2-yl)-2,5-diphenyl-2H-tetrazolium bromide (MTT) assay^71^; 200 µl fresh cell media and 250 µg of MTT was added to cells. Plates were Incubated at 37 °C, 5% CO_2_ for 3 hr then 200 µl of DMSO was added to wells with 25 µl glycine buffer (0.1 M glycine, 0.1 M NaCl pH 10.5) and shaken at 1000 r.p.m. for 10 min. Absorbance was read at OD_570_ nm. Cell viability was calculated as (sample absorbance – cell free blank/media control absorbance) x100.

## ACKNOWLEDGEMENTS

We are grateful to Magnus Unemo from WHO Collaborating Centre (CC) for Gonorrhoea and Other Sexually Transmitted Infections for providing the WHO reference gonococcal strains and to Eduard Sanders and his team at KEMRI institute in Kenya for providing the Kenyan clinical isolates. We would also like to thank Catherine Ison and Nerteley Quaye from the Sexually Transmitted Bacterial Reference Unit, Public Health England for UK *N. gonorrhoeae* clinical isolates and David Eyre from the Nuffield Department of Medicine, University of Oxford for *N. gonorrhoeae* G97687. We thank the University of Oxford’s Department of Chemistry Mass Spectrometry facility, especially Elisabete Pires and Jessica Pancholi, for their assistance with mass spectrometry. Furthermore, we would also like to thank Christopher Williams and Lianne Hill for mass spectrometry assistance from the University of Birmingham, The Centre for Chemical and Materials Analysis mass spectrometry facility.

## EXTENDED DATA

**Extended data table 1.** MtrE variants in *N. gonorrhoeae*.

| MtrE protein variant | MtrE (NEIS1632) alleles with the same a.a. sequence | Number of isolates | Percentage of isolates |
| --- | --- | --- | --- |
| 1 | 237, 238, 239, 286, 289, 300, 965, 1244, 1357, 1391, 1413, 1467 | 5863 | 31.4623 |
| 2 | 234, 247, 248, 283, 284, 285, 292, 295, 298, 365, 849, 906, 955, 1035, 1205, 1206, 1359, 1385, 1387, 1411, 1434, 1452 | 4043 | 21.6957 |
| 3 | 232, 240 241, 280, 287, 288, 971, 1389, 1453 | 3833 | 20.5688 |
| 4 | 235, 243, 293, 439, 919, 960, 967, 969, 1386, 1394, 1426, 1463 | 2276 | 12.2136 |
| 5 | 233 | 1902 | 10.2066 |
| 6 | 1570 | 282 | 1.5133 |
| 7 | 246, 281, 297 | 120 | 0.6439 |
| 8 | 1217, 1227, 1356, 1408 | 22 | 0.1181 |
| 9 | 302, 1448 | 21 | 0.1127 |
| 10 | 593 | 20 | 0.1073 |
| 11 | 1073 | 20 | 0.1073 |
| 12 | 236, 368, 839, 1010, 1409, 1433 | 18 | 0.0966 |
| 13 | 1237 | 17 | 0.0912 |
| 14 | 279 | 14 | 0.0751 |
| 15 | 330 | 13 | 0.0698 |
| 16 | 966 | 11 | 0.0590 |
| 17 | 818 | 9 | 0.0483 |
| 18 | 242 | 7 | 0.0376 |
| 19 | 304, 366, 1203 | 6 | 0.0322 |
| 20 | 838 | 6 | 0.0322 |
| 21 | 1031 | 6 | 0.0322 |
| 22 | 296 | 5 | 0.0268 |
| 23 | 1086 | 5 | 0.0268 |
| 24 | 1172 | 5 | 0.0268 |
| 25 | 1314 | 5 | 0.0268 |
| 26 | 920 | 4 | 0.0215 |
| 27 | 1465 | 4 | 0.0215 |
| 28 | 1468 | 4 | 0.0215 |
| 29 | 606 | 3 | 0.0161 |
| 30 | 610 | 3 | 0.0161 |
| 31 | 1084 | 3 | 0.0161 |
| 32 | 1208 | 3 | 0.0161 |
| 33 | 1435 | 3 | 0.0161 |
| 34 | 282 | 2 | 0.0107 |
| 35 | 290 | 2 | 0.0107 |
| 36 | 303, 1201 | 2 | 0.0107 |
| 37 | 592 | 2 | 0.0107 |
| 38 | 605 | 2 | 0.0107 |
| 39 | 611 | 2 | 0.0107 |
| 40 | 649 | 2 | 0.0107 |
| 41 | 817, 1245 | 2 | 0.0107 |
| 42 | 874 | 2 | 0.0107 |
| 43 | 959 | 2 | 0.0107 |
| 44 | 968 | 2 | 0.0107 |
| 45 | 1198 | 2 | 0.0107 |
| 46 | 1358 | 2 | 0.0107 |
| 47 | 1390 | 2 | 0.0107 |
| 48 | 1392 | 2 | 0.0107 |
| 49 | 1424 | 2 | 0.0107 |
| 50 | 1425 | 2 | 0.0107 |
| 51 | 1619 | 2 | 0.0107 |
| 52 | 245 | 1 | 0.0054 |
| 53 | 291 | 1 | 0.0054 |
| 54 | 294 | 1 | 0.0054 |
| 55 | 299 | 1 | 0.0054 |
| 56 | 301 | 1 | 0.0054 |
| 57 | 329 | 1 | 0.0054 |
| 58 | 367 | 1 | 0.0054 |
| 59 | 406 | 1 | 0.0054 |
| 60 | 428 | 1 | 0.0054 |
| 61 | 594 | 1 | 0.0054 |
| 62 | 607 | 1 | 0.0054 |
| 63 | 608 | 1 | 0.0054 |
| 64 | 609 | 1 | 0.0054 |
| 65 | 615 | 1 | 0.0054 |
| 66 | 715 | 1 | 0.0054 |
| 67 | 956 | 1 | 0.0054 |
| 68 | 957 | 1 | 0.0054 |
| 69 | 961 | 1 | 0.0054 |
| 70 | 1173 | 1 | 0.0054 |
| 71 | 1197 | 1 | 0.0054 |
| 72 | 1199 | 1 | 0.0054 |
| 73 | 1200 | 1 | 0.0054 |
| 74 | 1202 | 1 | 0.0054 |
| 75 | 1204 | 1 | 0.0054 |
| 76 | 1207 | 1 | 0.0054 |
| 77 | 1209 | 1 | 0.0054 |
| 78 | 1218 | 1 | 0.0054 |
| 79 | 1228 | 1 | 0.0054 |
| 80 | 1229 | 1 | 0.0054 |
| 81 | 1230 | 1 | 0.0054 |
| 82 | 1250 | 1 | 0.0054 |
| 83 | 1360 | 1 | 0.0054 |
| 84 | 1361 | 1 | 0.0054 |
| 85 | 1362 | 1 | 0.0054 |
| 86 | 1384 | 1 | 0.0054 |
| 87 | 1388 | 1 | 0.0054 |
| 88 | 1393 | 1 | 0.0054 |
| 89 | 1395 | 1 | 0.0054 |
| 90 | 1410 | 1 | 0.0054 |
| 91 | 1412 | 1 | 0.0054 |
| 92 | 1464 | 1 | 0.0054 |
| 93 | 1466 | 1 | 0.0054 |
| 94 | 1577 | 1 | 0.0054 |

**Extended data table 2.**
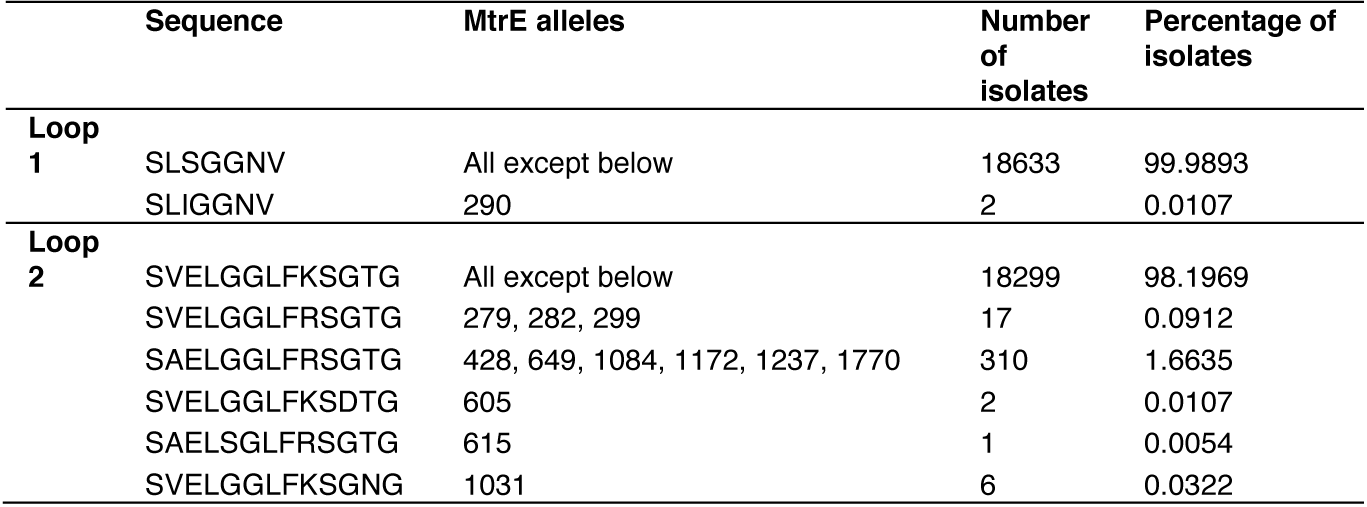
*N. gonorrhoeae* MtrE outer membrane loop sequences.

|  |  |  |  |  |
| --- | --- | --- | --- | --- |
|  | Sequence | MtrE alleles | Number of isolates | Percentage of isolates |
| Loop 1 | SLSGGNV | All except below | 18633 | 99.9893 |
|  | SLIGGNV | 290 | 2 | 0.0107 |
| Loop 2 | SVELGGLFKSGTG | All except below | 18299 | 98.1969 |
|  | SVELGGLFRSGTG | 279, 282, 299 | 17 | 0.0912 |
|  | SAELGGLFRSGTG | 428, 649, 1084, 1172, 1237, 1770 | 310 | 1.6635 |
|  | SVELGGLFKSDTG | 605 | 2 | 0.0107 |
|  | SAELSGLFRSGTG | 615 | 1 | 0.0054 |
|  | SVELGGLFKSGNG | 1031 | 6 | 0.0322 |

**Extended data table 3a.** *Neisseria meningitidis* and commensal *Neisseria* spp. used in this study.

| PubMLST ID | Species (*Neisseria*) | Isolate | Country | Year | MtrE (NEIS1632) allele | Reference |
| --- | --- | --- | --- | --- | --- | --- |
| 698 | *meningitidis* | Fam18 | USA | 1983 | 1 | Feavers et al. (1999) |
| 1038 | *meningitidis* | 8013 | France | 1989 | 16 | Rusniok et al. (2009) |
| 240 | *meningitidis* | MC58 | UK | 1983 | 3 | Tettelin et al. (2005) |
| 34542 | *meningitidis* | NZ 98/254 | New Zealand | 1998 | 19 | Findlow et al. (2006) |
| 1773 | *lactamica* | 042-12 | UK | 1998 | - | Bennett et al. (2005) |
| 1775 | *lactamica* | 047-24 | UK | 1999 | - | Bennett et al. (2005) |
| 1781 | *lactamica* | 004-24 | UK | 1998 | - | Bennett et al. (2005) |
| 1793 | *lactamica* | 005-24 | UK | 1998 | - | Bennett et al. (2005) |
| 8851 | *lactamica* | 020-06 | UK | 1997 | 4 | Bennett et al. (2005) |
| 49342 | *cinerea* | 5746 | Sweden | 1977 | 1826 | Bennett et al. (2012) |
| 49343 | *cinerea* | CCUG 25879 | Sweden | 1989 | 1827 | Bennett et al. (2012) |
| 49344 | *cinerea* | CCUG 27178A | Unknown | 1983 | 1828 | Bennett et al. (2012) |
| 21041 | *cinerea* | CCUG 53043 | Sweden | 2006 | 1814 | Bennett et al. (2013) |
| 49341 | *cinerea* | CCUG 346T | Unknown | Unknown | 1825 | Bennett et al. (2012) |
| 49356 | *polysaccharea* | CCUG 18031 | Germany | 1984 | 785 | Bennett et al. (2012) |
| 49358 | *polysaccharea* | CCUG 24846 | Unknown | Unknown | 1875 | Bennett et al. (2012) |
| 49359 | *polysaccharea* | CCUG 27182 | Unknown | Unknown | 76 | Bennett et al. (2012) |
| 49357 | *polysaccharea* | CCUG 24845 | Unknown | Unknown | 80 | Bennett et al. (2012) |
| 49339 | *bergeri* | 15883 | Germany | 1984 | 1873 | Bennett et al. (2012) |

**Extended data table 3b.** Presence or absence of IgAP in *Neisseria meningitidis* and commensal *Neisseria* spp. used in this study.

| Species | Presence of NEIS0651 (IgAP) |
| --- | --- |
| *Neisseria gonorrhoeae* | 19,865 / 19,865 |
| *Neisseria lactamica* | 0 / 1118 |
| *Neisseria polysaccharea* | 0 / 71 |
| *Neisseria cinerea* | 0 / 16 |
| *Neisseria subflava* | 0 / 71 |
| *Neisseria bergeri* | 0 / 61 |
| *Neisseria mucosa* | 0 / 25 |

**Extended data 1.**
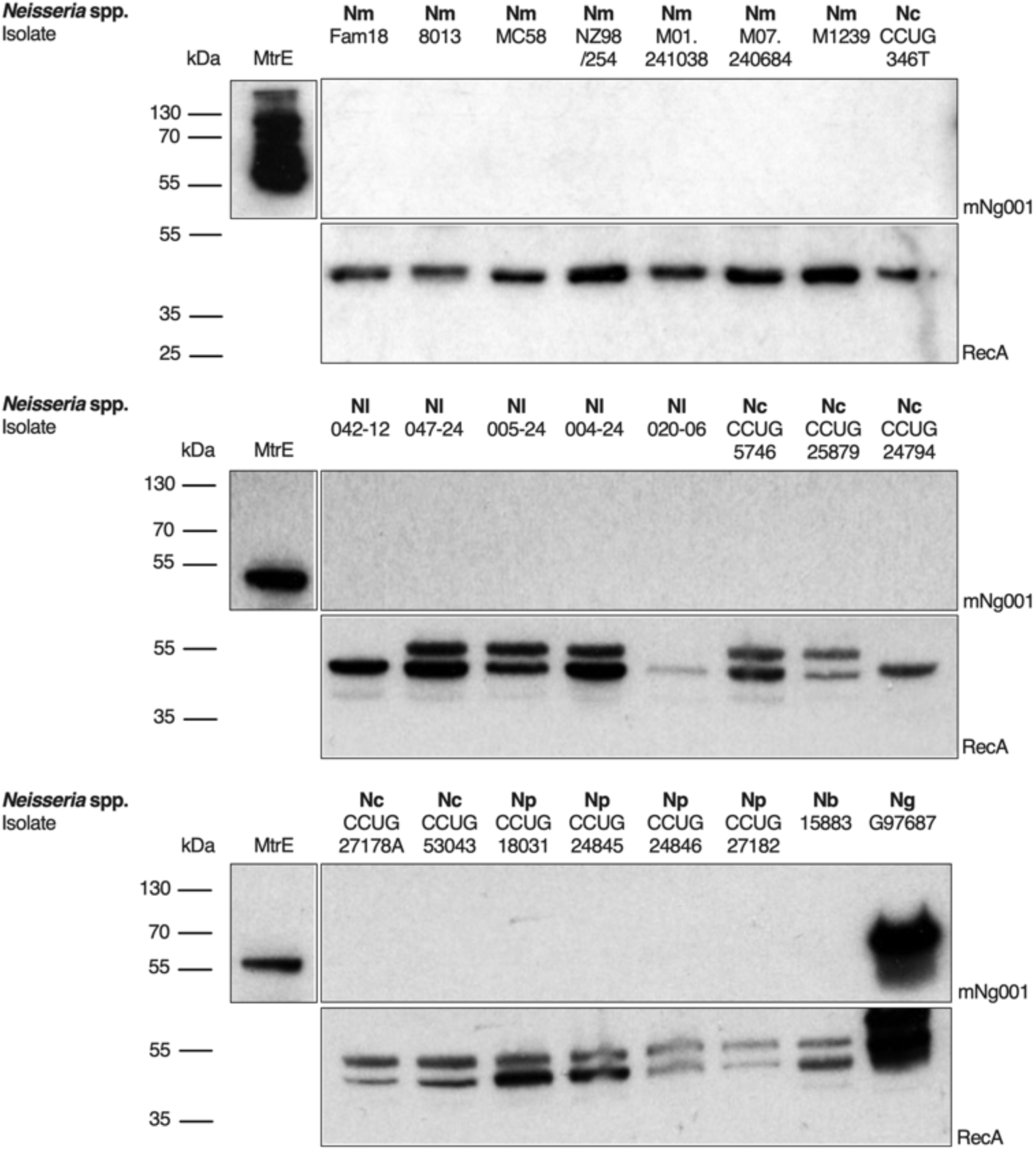
Monoclonal antibody, mNg001 does not recognise MtrE from *Neisseria meningitidis* or non-invasive *Neisseria* spp. Western blot analysis for recognition of MtrE from; *N. meningitidis* (Nm), *Neisseria lactamica* (Nl), *Neisseria cinerea* (Nc), *Neisseria polysaccharea* (Np) and *Neisseria bergeri* (Nb) using mAb, mNg001. Recombinant MtrE and wild-type MDR gonococcal strain (Ng), G97687 were used as positive controls; RecA used as a loading control.

**Extended data 2.**
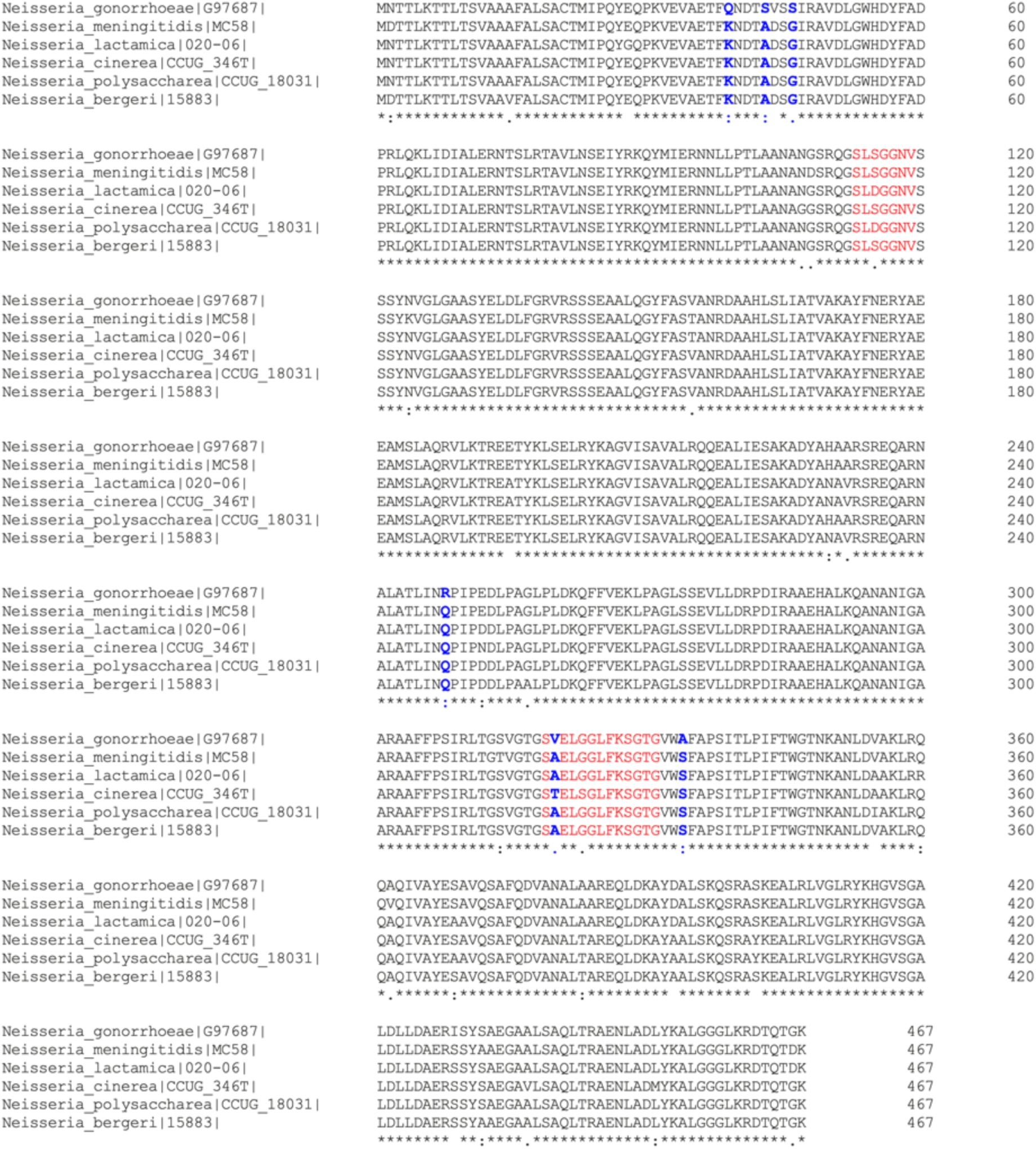
MtrE from *N. gonorrhoeae* shows consistent sequence differences compared to other *Neisseri*a spp. Sequence alignment of MtrE from a representative strain from different Neisseria spp. compared to MtrE from *N. gonorrhoeae* strain G97687, the most common gonococcal MtrE protein variant. Red residues indicate the two surface exposed loops whereas blue residues indicate consistent sequence differences from gonococcal MtrE.

**Extended data 3.**
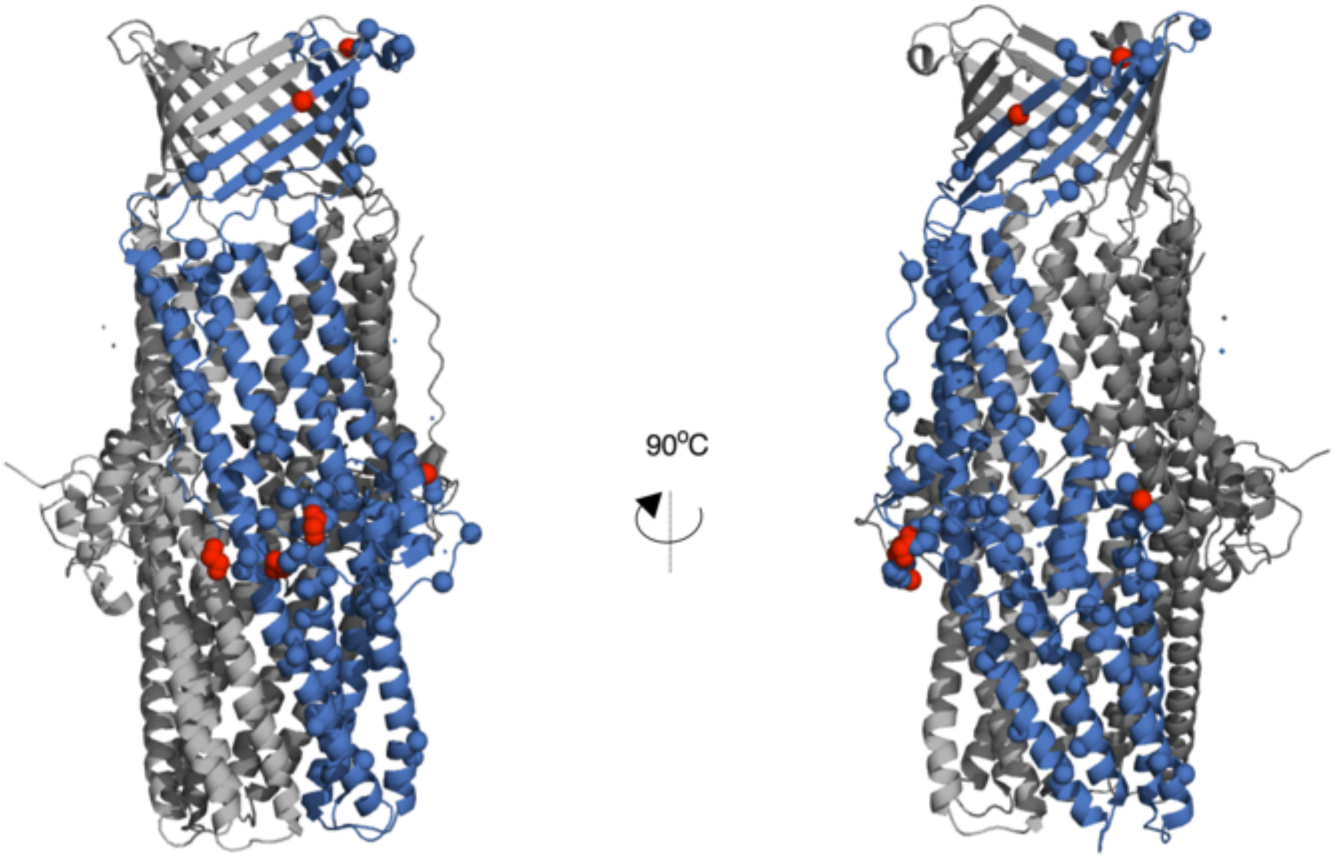
Polymorphisms in MtrE from *N. meningitidis* and non-invasive *Neisseria* spp. Red amino acids indicate polymorphisms which are different between the five most common gonococcal MtrE variants and MtrE found in *N. meningitidis* and non-invasive *Neisseria* mapped onto trimeric MtrE (PDB accession number 4MT0); MtrE monomer (blue)

**Extended data 4.**
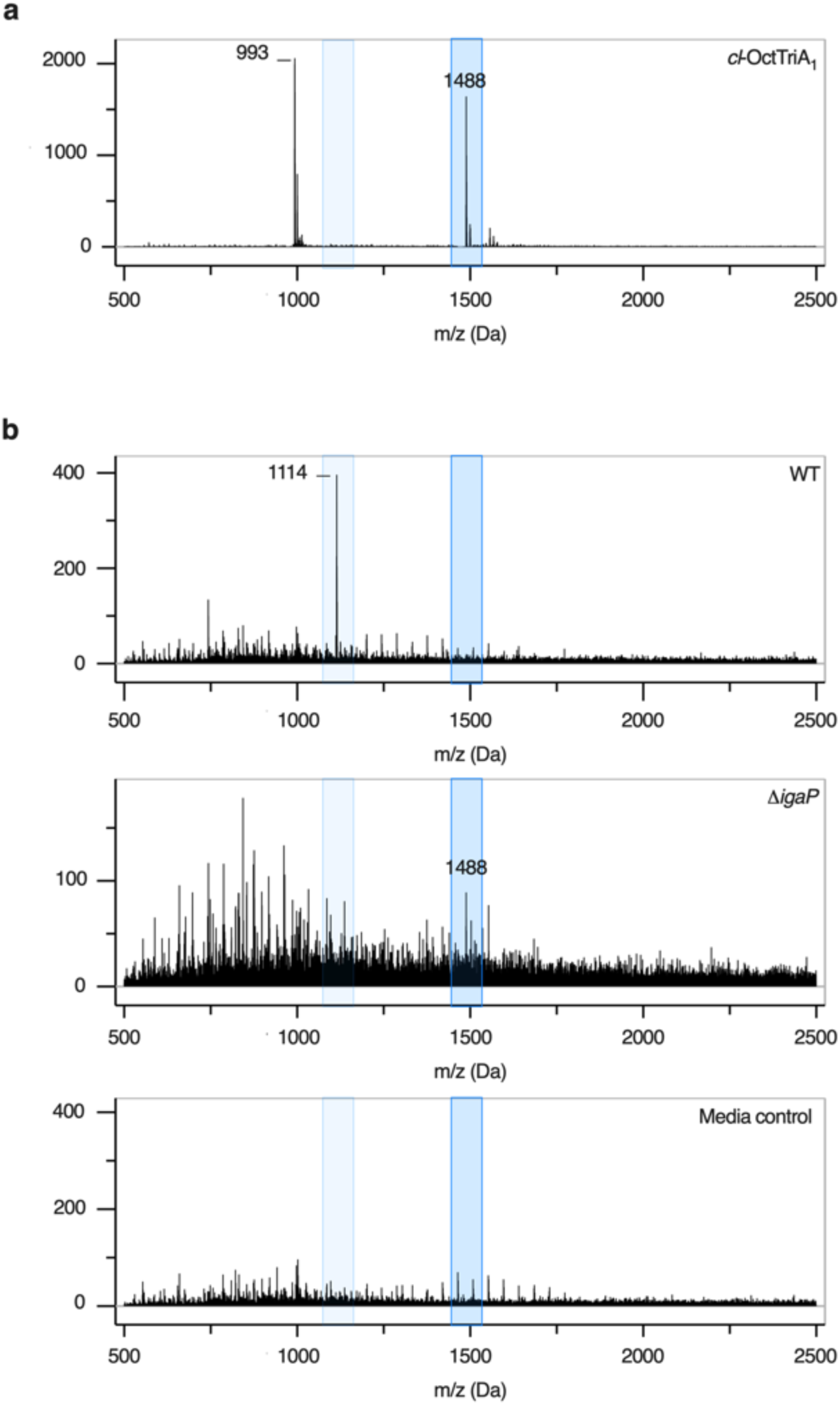
Peptide *cl*-OctTriA_1_ is specifically cleaved by the gonococcal IgAP. **(a)** Synthesised *cl*-OctTriA_1_-*c* (dark blue box) in PBS shown as a mass spectrometry control. **(b)** Analysis of *cl*-OctTriA_1_ after incubation in supernatants from wild-type FA1090 (top) but not FA1090Δ*igaP* (middle) by Electron spray isonisation mass spectrometry. MS species corresponding to peptide *cl*-OctTriA_1_ (dark blue box) and the cleaved product OctTriA_1_-*c* (light blue box) are indicated in each spectrum. Media only control to indicate background (bottom).

**Extended data 5.**
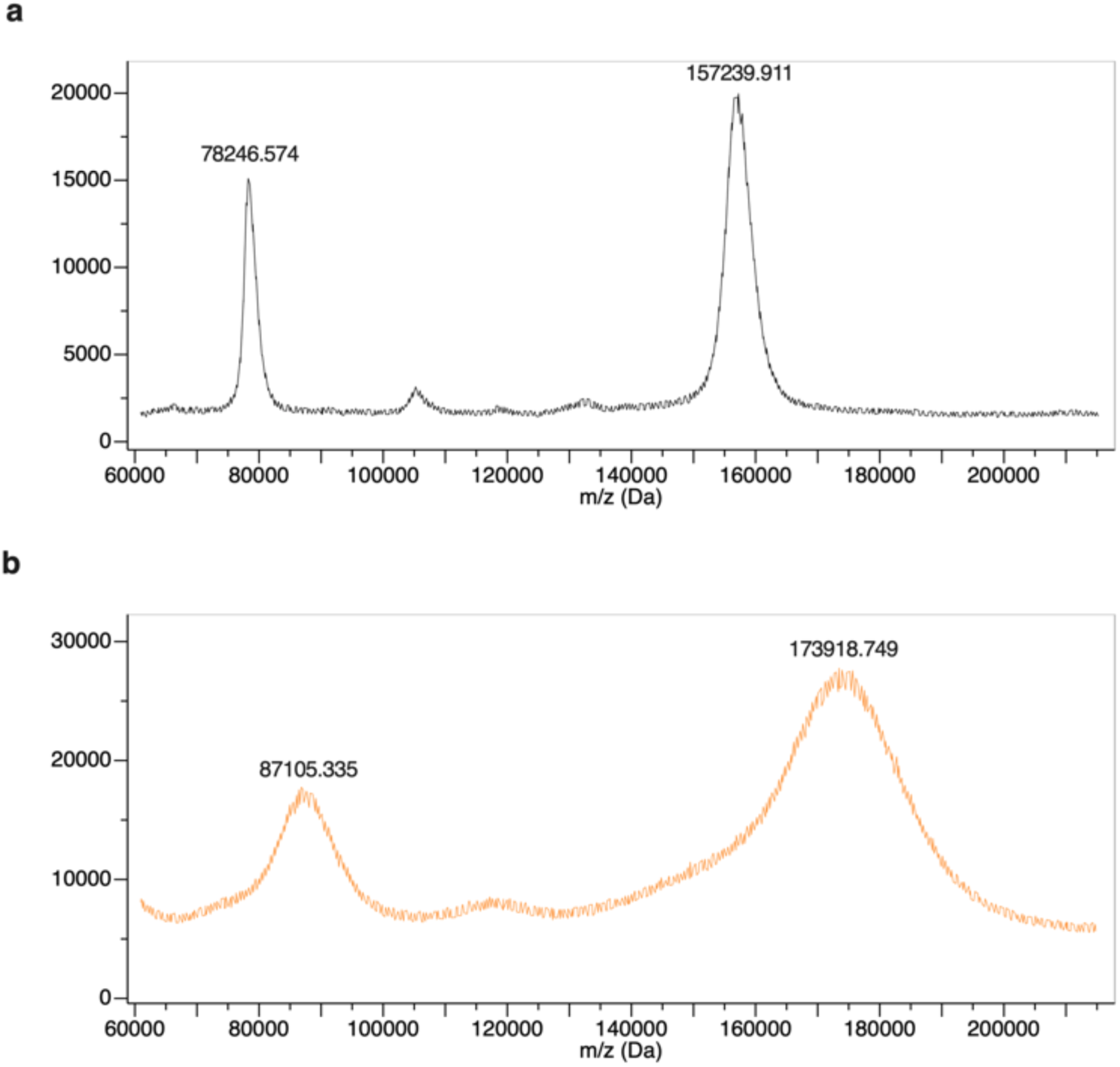
Non-cleavable ADCs have a similar DAR to IgAP cleavable ADCs. MALDI-TOF analysis of mNg001 (top, black) and ADC covalently linked to synthesized uncleavable OctTriA_1_ analogue, uncl-Oct-TriA_1_ (bottom, orange). Change in mass, compared to mNg001 indicates a DAR of approximately 6 peptides per ADC.

**Extended data 6.**
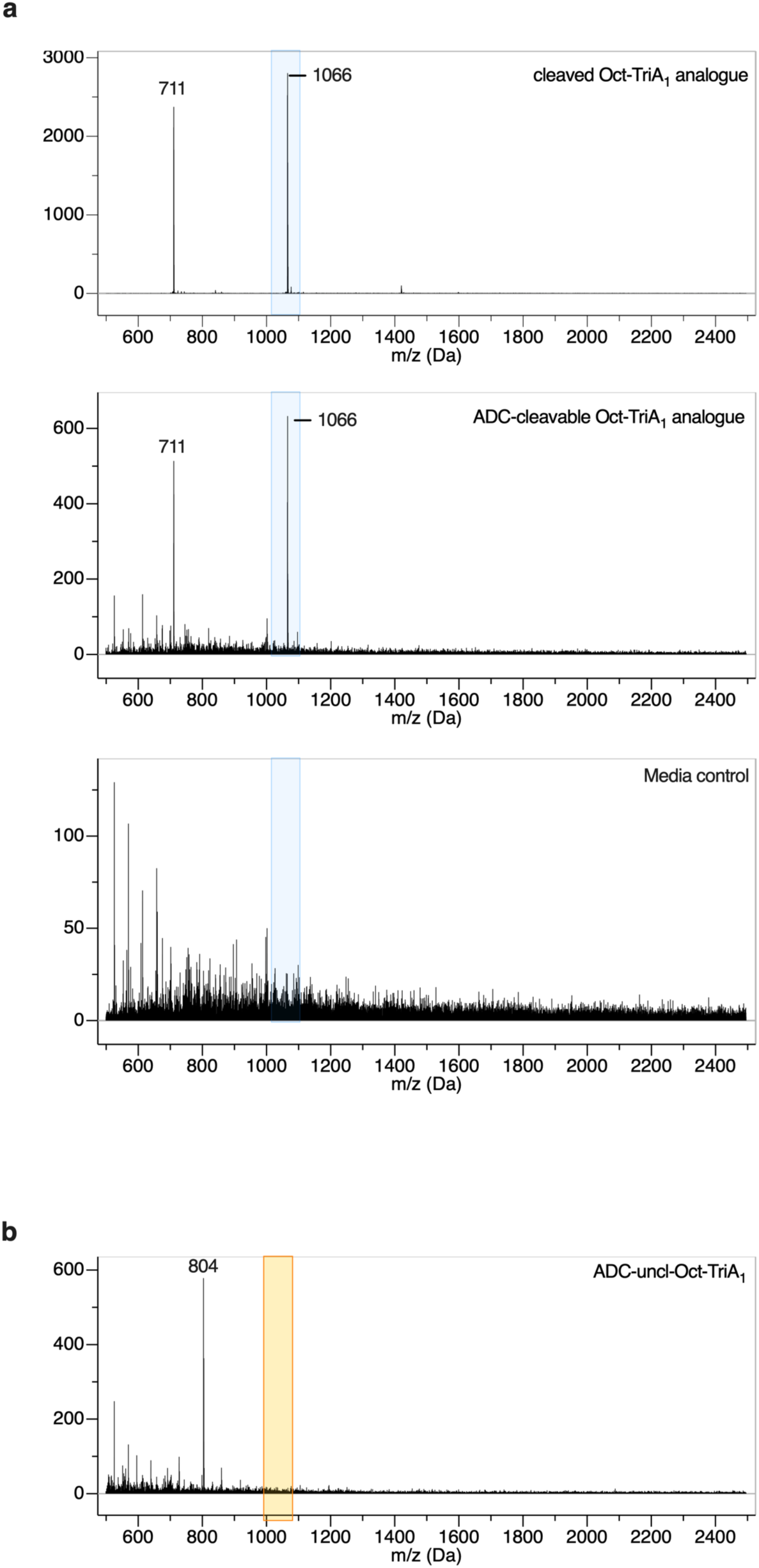
Peptide *uncl*-OctTriA_1_ is not cleaved from the ADC by the gonococcal IgAP. **(a)** Synthesised cleaved OctTriA_1_ (cleaved linker sequence Val-Val-Ala-Pro-Pro) (blue box, top) in PBS shown as a mass spectrometry control. Analysis of cleavage of OctTriA_1_ from ADC after incubation in supernatant from wild-type FA1090 (middle) by Electron spray isonisation mass spectrometry. Media only control to indicate background (bottom). **(b)** No peptide release was observed after incubation of ADC-uncl-OctTriA_1_ with supernatant from FA1090 (orange box).

**Extended data 7.**
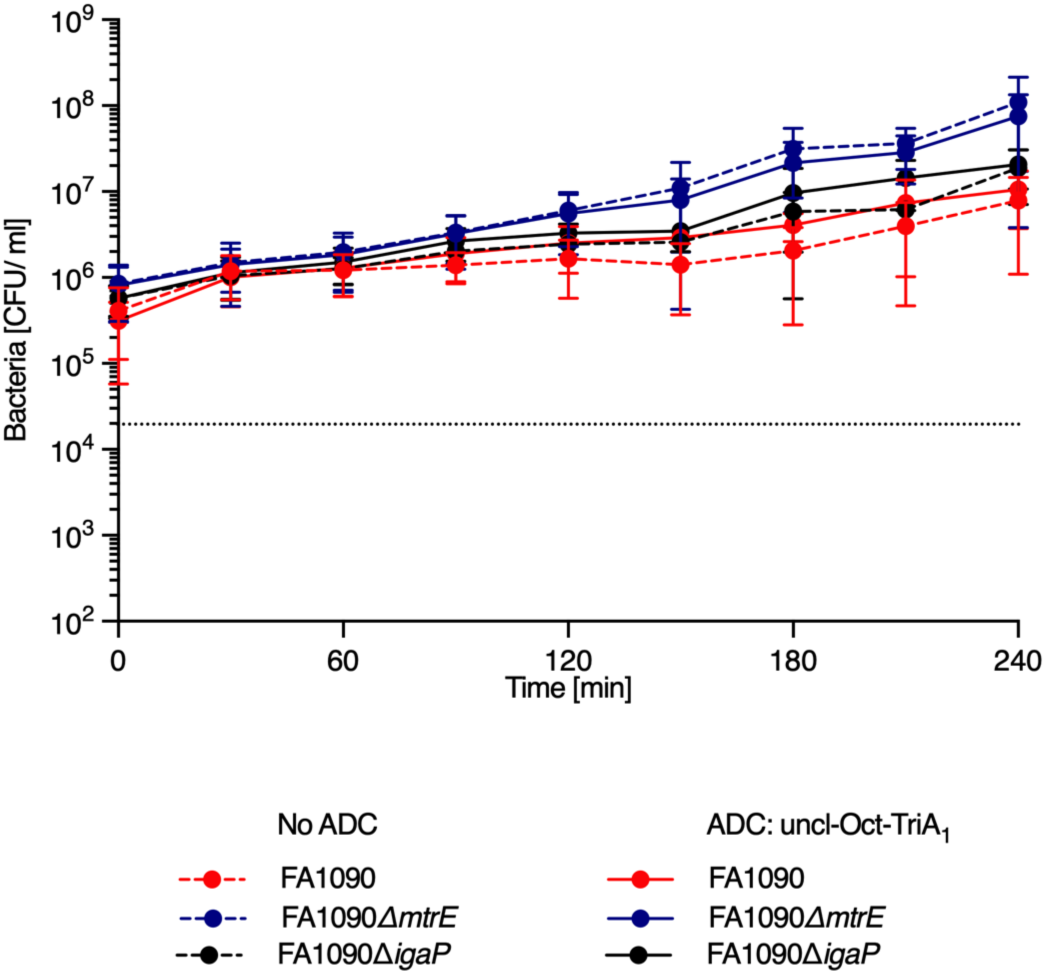
Non-cleavable ADCs are not bactericidal in the presence of the gonococcal IgAP. Time to kill curve analysis of ADC-un*cl*-OctTriA_1_ against FA1090 (red), isogenic *ΔigAP* (black) and *ΔmtrE* (blue). No ADC (dashed line) and ADC (line). All data shown are the mean ± standard deviation of CFU/ml (n = 3).

## SUPPLEMENTARY DATA

**Supplementary table 1.** Oct-TriA1 analogue sequences used in this study.

| Peptide name | Peptide | Peptide sequence |
| --- | --- | --- |
| Oct-TriA<sub>1</sub> | Oct-TriA<sub>1</sub> | Octanoyl-DVal-DDab-Gly-DSer-DTrp-Ser-Dab-DDab-Phe-Glu-Val-Dalle-Ala-OH |
| d-Oct-TriA<sub>1</sub> | Oct-TriA<sub>1</sub>-(C) | Octanoyl-DVal-DDab-Gly-DSer-DTrp-Ser-Dab-DDab-Phe-Glu-Val-Dalle-Ala-O2Oc-O2Oc-Cys-NH2 |
| Oct-TriA<sub>1</sub>-c | Oct-TriA<sub>1</sub>-(LPRPP) | Octanoyl-DVal-DDab-Gly-DSer-DTrp-Ser-Dab-DDab-Phe-Glu-Val-Dalle-Ala-O2Oc-Leu-Pro-Arg-Pro-Pro-OH |
| cl-Oct-TriA<sub>1</sub> | Oct-TriA<sub>1</sub>-(LPRPPAPVFS-C) | Octanoyl-DVal-DDab-Gly-DSer-DTrp-Ser-Dab-DDab-Phe-Glu-Val-Dalle-Ala-O2Oc-Leu-Pro-Arg-Pro-Pro-Ala-Pro-Val-Phe-Ser-O2Oc-Cys-NH2 |
| uncl-Oct-TriA<sub>1</sub> | Oct-TriA<sub>1</sub>-(VKPAPSPAAN-C) | Octanoyl-DVal-DDab-Gly-DSer-DTrp-Ser-Dab-DDab-Phe-Glu-Val-Dalle-Ala-O2Oc-Val-Lys-Pro-Ala-Pro-Ser-Pro-Ala-Ala-Asn-O2Oc-Cys-NH2 |

**Supplementary table 2.**
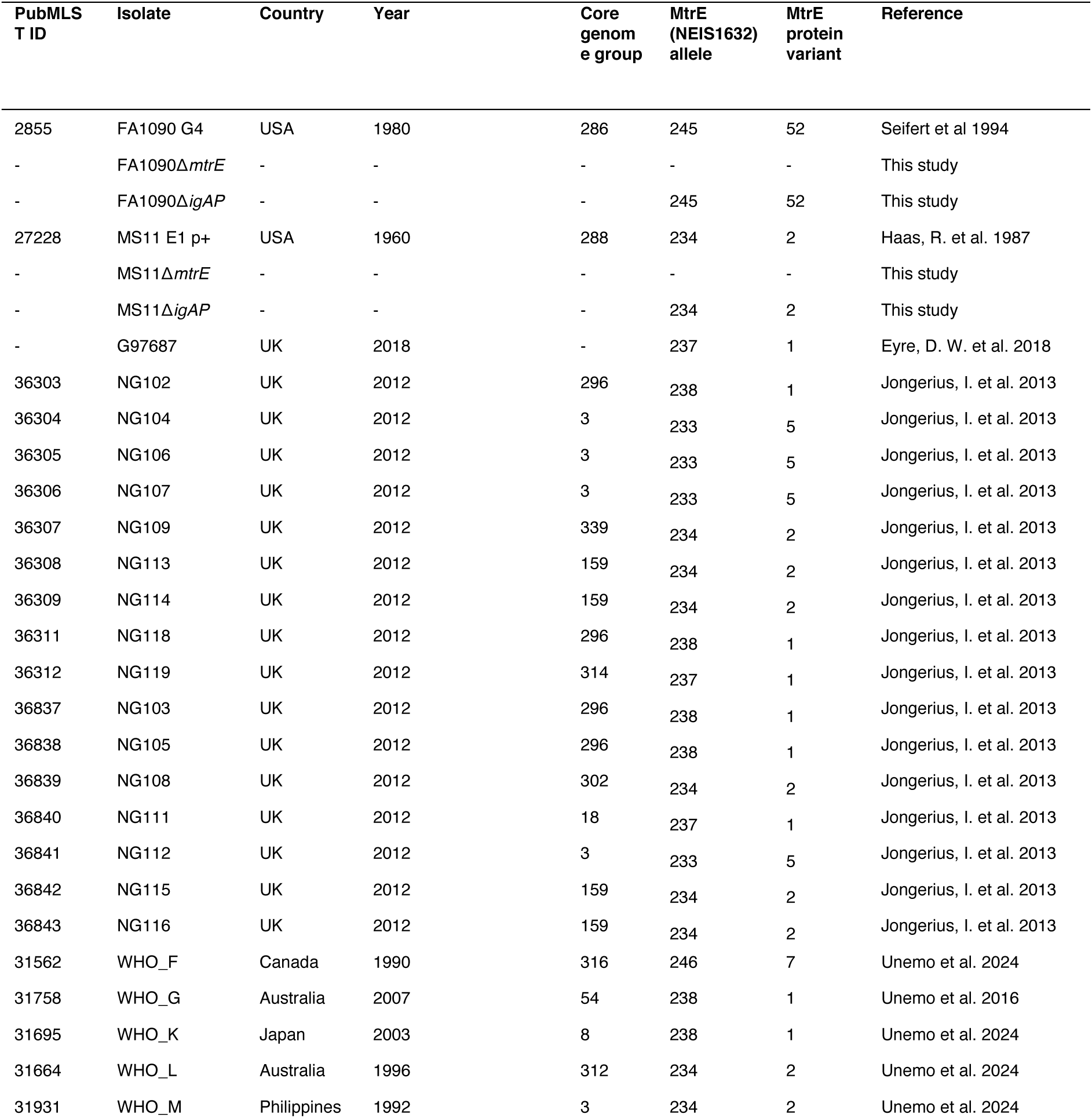
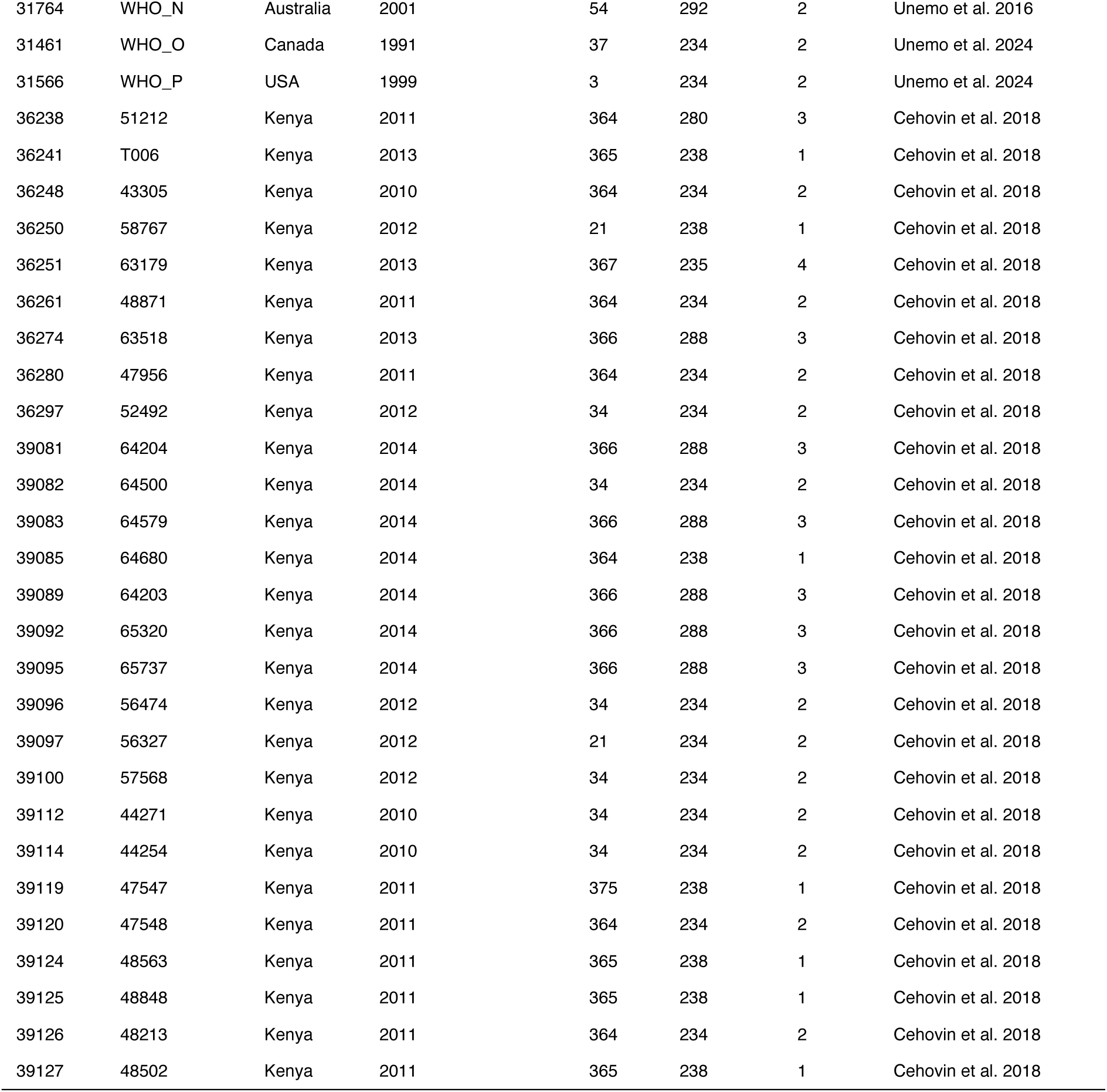
*N. gonorrhoeae* strains used in this study.

**Supplementary table 3.**
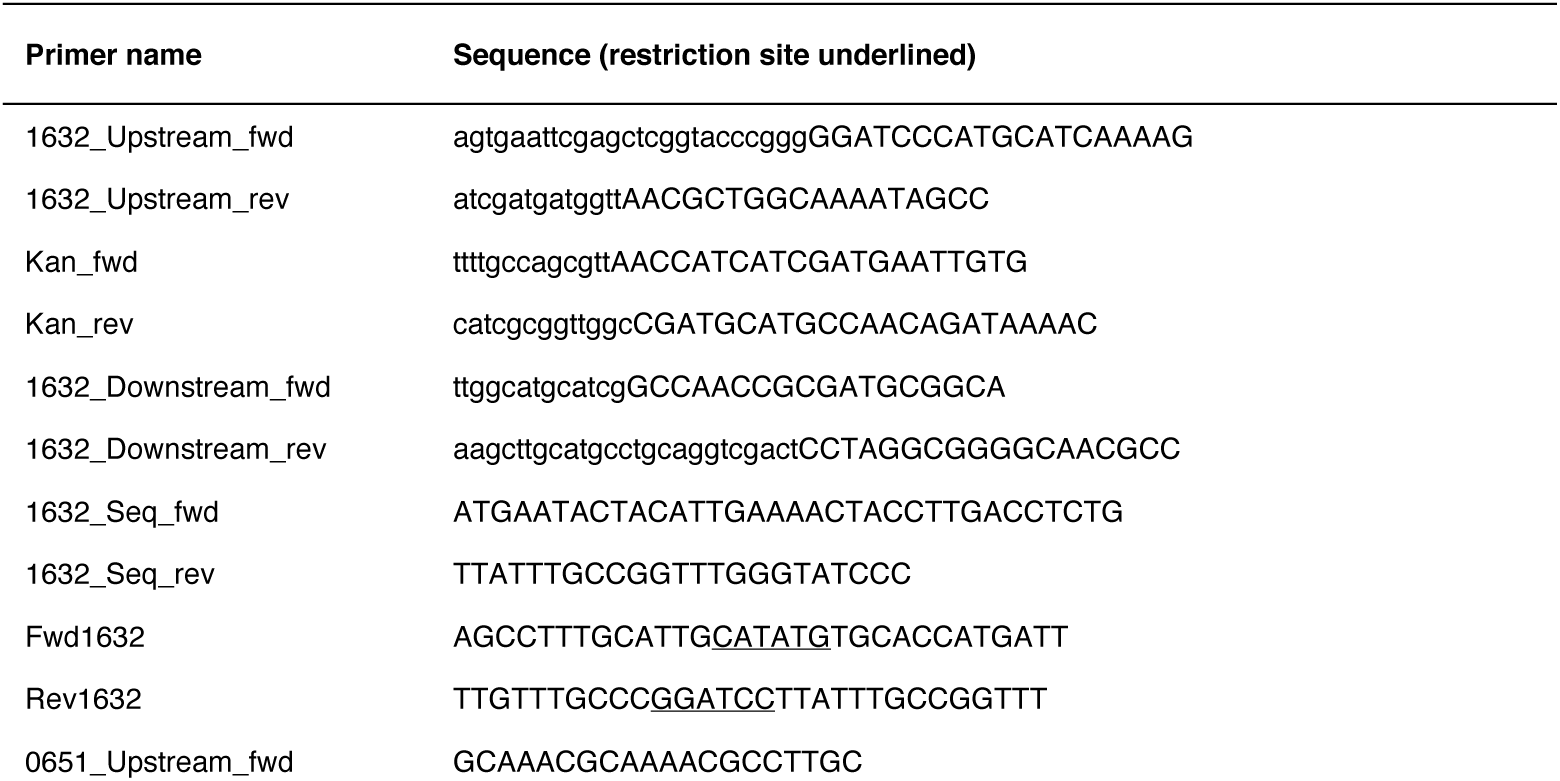
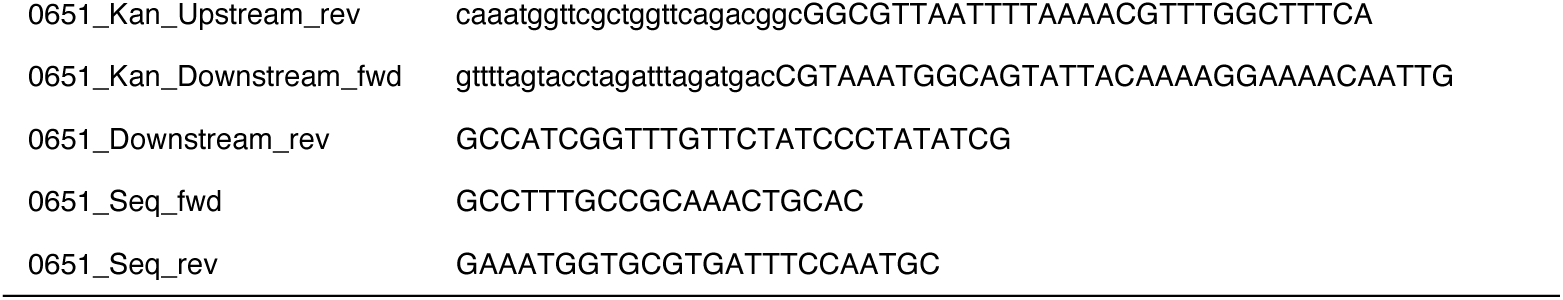
Primers used in this study.

